# Decoding spatiotemporal gene expression of the developing human spinal cord and implications for ependymoma origin

**DOI:** 10.1101/2022.08.31.505986

**Authors:** Xiaofei Li, Zaneta Andrusivova, Paulo Czarnewski, Christoffer Mattsson Langseth, Alma Andersson, Yang Liu, Daniel Gyllborg, Emelie Braun, Ludvig Larsson, Lijuan Hu, Zhanna Alekseenko, Hower Lee, Christophe Avenel, Helena Kopp Kallner, Elisabet Åkesson, Igor Adameyko, Mats Nilsson, Sten Linnarsson, Joakim Lundeberg, Erik Sundström

## Abstract

The human spinal cord contains diverse cell types, governed by a series of spatiotemporal events for tissue assembly and functions. However, the spatiotemporal regulation of cell fate specification in the human developing spinal cord remains largely unknown. Single-cell RNA sequencing and spatial transcriptomics techniques have advanced the understanding of human organ development considerably. By performing integrated analysis of single-cell and spatial multi-omics methods, we created a comprehensive developmental cell atlas of the first trimester human spinal cord. Our data revealed that the cell fate commitment of neural progenitor cells and their spatial positioning are spatiotemporally regulated by specific gene sets. Beyond this resource, we unexpectedly discovered unique events in human spinal cord development compared to rodents, including earlier quiescence of active neural stem cells, different regulation of stem cell differentiation, and distinct spatiotemporal genetic regulations of cell fate choices. In addition, using our atlas we identified specific gene expression in cancer stem cells in ependymomas. Thus, we demonstrate spatiotemporal genetic regulation of human spinal cord development as well as its potential to understand novel disease mechanisms and to inspire new therapies.

## Main

The spinal cord comprises the caudal region of the central nervous system (CNS) and is responsible for conveying and processing motor and sensory information between the brain and the periphery, as well as for elaborating reflexes. During spinal cord development, neural stem and progenitor cells (NPCs) in the ventricular zone, surrounding the nascent central canal, are committed to their respective cell fates governed by gradients of dorsal and ventral morphogens^1^. Consequently, different transcription factors (TFs) along the dorsal-ventral (DV) axis are activated, resulting in spatially segregated progenitor domains. In rodents, domain-specific NPCs temporally undergo cell fate specification, first generating neurons, then glia. These differentiated neural cells migrate from their origin to their final locations in the spinal cord and engage in distinct circuits ^1^.

It is however, not known to what extent this knowledge can be translated to humans. It is generally believed that during the first trimester of human pregnancy, most of the human (h)NPCs are highly proliferative in preparation for neurogenesis and gliogenesis. Therefore, cell therapy approaches such as stem cell therapies for neurotrauma and degenerative diseases, usually obtain hNPCs from the first trimester, thus more likely to acquire active and neuron-fate-committed NPCs. Current studies, however, showed that hNPCs derived from early development exhibit either robust glial differentiation^2^ or little differentiation ^3^, suggesting that besides the impacts of microenvironment, deciphering the intrinsic genetic regulation for cell fate commitment of hNPCs is necessary to achieve better efficiency of such therapies. Furthermore, impaired neurodevelopment, pediatric tumorigenesis and neurodevelopmental disorders are highly related. Therefore, better understanding of the cell fate commitment of hNPCs in the developing spinal cord can provide insights into human developmental biology, future regenerative strategies and potential pediatric cancer treatment.

Single-cell RNA sequencing (scRNA-seq) and spatial transcriptomics (ST) have provided high-throughput and spatially resolved analysis of spatiotemporal gene expression during human prenatal development ^4^. Furthermore, a high-throughput and multiplex *in situ* hybridization method, hybridization-based *in situ* sequencing (HybISS) has recently been developed for single RNA molecule localization of large gene panels with single-cell resolution within human tissue for data validation ^4, 5^. Combining all these methods can largely reduce the limitations of individual techniques, facilitate unbiased cell type annotation, and allow high resolution spatiotemporal mapping of the developing human spinal cord. Two recent studies used scRNA-seq on human developing spinal cord and revealed the appearance of different neural cell types ^6, 7^. However, the genetic regulation of the commitment of homogenous hNPCs to heterogenous neuronal and glial fates *in vivo* is still unclear. Furthermore, although neural patterning associated with transient spatial distribution of neural cells during human development is well-known ^7, 8^, how such events result in spatially restricted heterogenous neurons and glia is still not well studied in human. In addition, while the use of transgenic animals during the last two decades has provided detailed mechanisms of the rodent spinal cord development, it is unclear whether there are unique features of the development of the human spinal cord. In this study, we have analyzed human embryonic and fetal spinal cords covering the entire first trimester, using state-of-the-art scRNA-seq, ST and HybISS, and integrated these datasets with previously reported mouse and human spinal cord datasets for our analysis. Here we provide a comprehensive developmental cell atlas of the human spinal cord, reveal spatiotemporal gene expression and regulation of cell fate commitment, highlight the major differences of cellular and molecular events in human and rodent spinal cord development, and discover novel molecular targets and genetic regulation of pediatric spinal cancer stem cells.

### Comprehensive atlas of the human developing spinal cord

To investigate the molecular features of the developing human spinal cord, we acquired 16 human prenatal spinal cords at post-conception week (W) 5-12 (Supplementary Table 1), covering the first trimester of pregnancy when cell fate specifications in the CNS occur ^9, 10^. We performed scRNA-seq, ST and HybISS to create a developmental cell atlas of the human spinal cord with detailed spatiotemporal gene expression and validation (Fig. 1a, Supplementary Table 1). A total of 159,350 high quality cells across 31 scRNA-seq libraries were analyzed, revealing 47 cell clusters (Extended Data Fig. 1a-b) (16 major cell populations) (Fig. 1b). All major spinal cord neural cell types were represented including NPCs, intermediate neuronal progenitors (INPs), excitatory neurons (ExNs), inhibitory neurons (IbNs), cholinergic neurons (ChNs), astrocytes (ASCs), ependymal cells (EPCs), oligodendrocyte precursor cells (OPCs) and oligodendrocytes (OLs) (Fig. 1b), which this study mainly focused on. Other cell types such as Schwann cells (SWCs), pericytes (PCs), endothelial cells (ENs), vascular capillary endothelial cells (VCLPs) and immune cells (Immune) (e.g., microglia) were also derived during this developmental stage (Fig. 1b). Top marker genes of each cell type and cluster are summarized (Fig. 1c, Extended Data Fig. 1c).

**Fig. 1:**
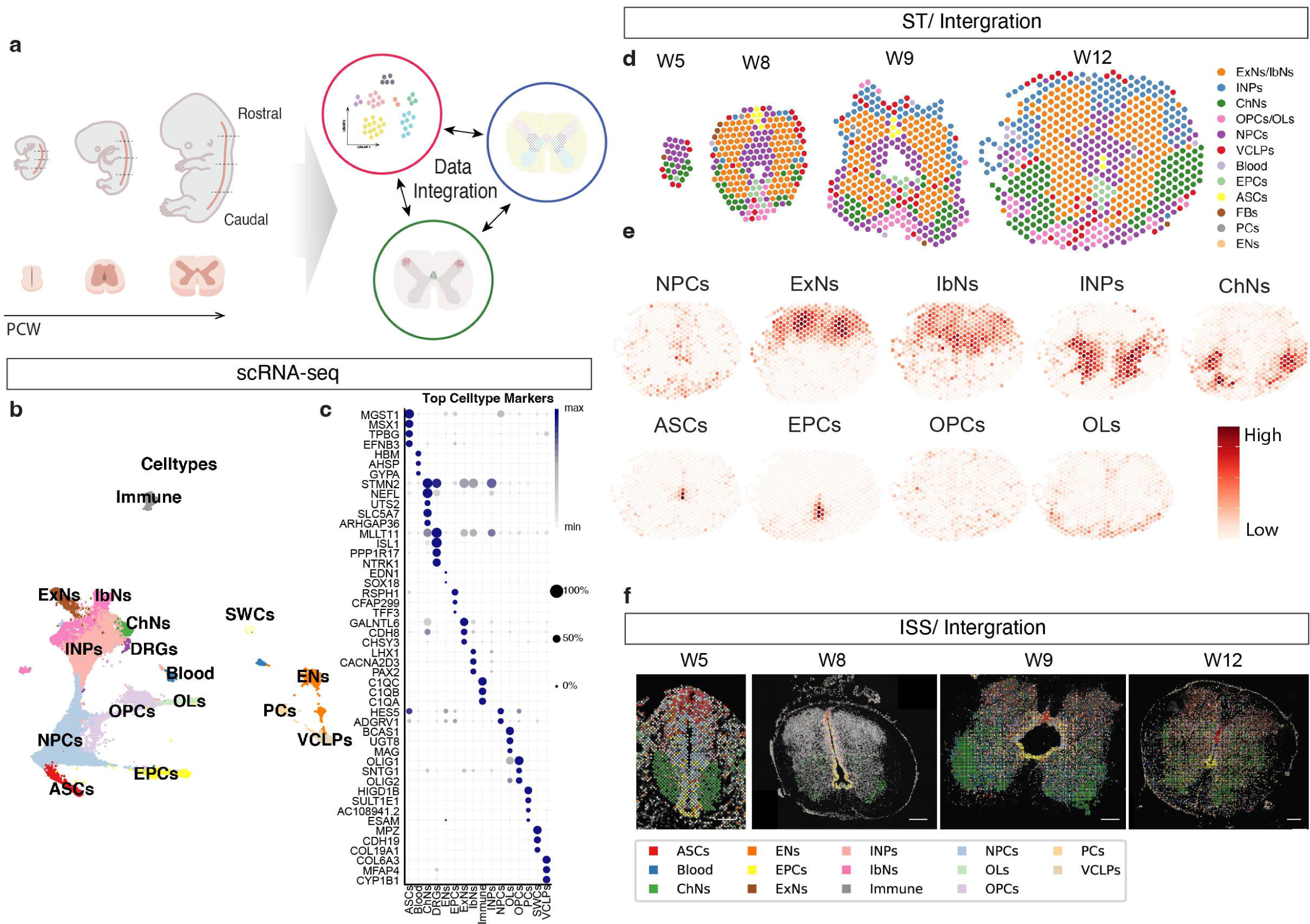
Comprehensive atlas of the developing human spinal cord. a) Schematic overview of the workflow. b) UMAP of scRNA-seq datasets revealing major cell populations. c) Dot plot illustrating top marker genes for major cell populations. d) Spatial mapping of major cell types from ST analysis in representative human spinal cord sections. e) Representative stereoscope plots of one W12 section. f) Representative images and cell typing results from HybISS. Scale bar = 200μm. Neural stem and progenitor cells (NPCs), intermediate neuronal progenitors (INPs), excitatory neurons (ExNs), inhibitory neurons (IbNs), cholinergic neurons (ChNs), astrocytes (ASCs), ependymal cells (EPCs), oligodendrocyte precursor cells (OPCs), oligodendrocytes (OLs), immune cells (Immune), Schwann cells (SWCs), pericytes (PCs), endothelial cells (ENs) and vascular capillary endothelial cells (VCLPs).

To define the spatial gene expression and analyze cell type localization independently from scRNA-seq, we used sections of the prenatal spinal cords along the rostral-caudal axis (RC) of representative ages (W5, W8, W9 and W12) for analysis. Clustering analysis of ST data from 76 sections resulted in 23 clusters along the RC and DV axis (Extended Data Fig. 2a-b,d), and revealed 12 major cell types (Fig. 1d, Extended Data Fig. 2c). At W5, the cross-sectioned human spinal cord was dominated by NPCs in the ventricular zone. From W8 and onwards, not only neurons but all glial cell types were born (Fig. 1d, Extended Data Fig. 2c). In addition, fewer cell types could be identified in the caudal regions with our clustering approach (e.g., cluster 0 neurons at W8) compared to the rostral regions, suggesting a possible earlier development in rostral regions (Extended Data Fig. 2c and e). However, we did not obtain obvious differences in gene expression by comparing scRNA-seq data and ST data from different regions along the RC axis, suggesting that the asymmetric development along RC axis could be regulated by secreted factors at protein level. To understand the probability of cell types in different of the human spinal cord, we further integrated scRNA-seq and ST data by using *stereoscope*, a recently developed method for guided decomposition of ST data by using scRNA-seq data as reference ^11^ to delineate the spatial distribution of cell types defined in the scRNA-seq. (Fig. 1e, Extended Data Fig. 3). We found that the dorsal area was mainly occupied by ExNs and IbNs, while the ventral gray matter was mainly occupied by immature neurons and ChNs from W8 (Fig. 1e). Early born glial cells showed cell type specific spatial distributions, including ASCs in the dorsal ventricular zone and EPCs and OPCs in the ventral ventricular zone (W5 and W8 data in Extended Data Fig. 3). Notably, stereoscope data indicates the relative probability of each cell type in certain spot, rather than an absolute value of cell number quantification. To provide single cell spatial mapping resolution and validation, we performed HybISS ^5^ in adjacent tissue sections to visualize the transcriptome *in situ* using 50 selected genes (Supplementary Table 3; Supplementary Figure 1) for major cell type characterization and 224 genes for subtype or cell state characterization (Supplementary Table 4; Supplementary Figure 2). The HybISS data were integrated with scRNA-seq data by probabilistic cell typing (pciSeq) ^12^, and confirmed the findings revealed by ST (Fig. 1f). Notably, either ST or HybISS was also analyzed independently from scRNA-seq data as validation (more detail below).

### Heterogenous neural cells in the human developing spinal cord

To validate the major cell populations that we identified with scRNA-seq and ST (Fig. 1 b,d,e), we selected 50 genes matching the markers of each major cell type (Supplementary Figure 1) and performed HybISS on the human spinal cord sections. We observed that in agreement with ST and stereoscope data (Fig. 1d-e), NPCs (*ASCL1+ SOX2+*) were the major cell population at W5 and were highly proliferative (*MKI67+TOP2A+*), but from W8 the proliferating NPCs were restricted to the ventricular zone (Fig 2a). Different neurons, including ExNs (e.g. *CACN2D1+*), IbNs (e.g. *SCGZ+* or *NRXN3+*) and ChNs (*ISL1*+ and/or *SLC5A7*+) appeared as early as W5 and were widely distributed throughout the gray matter (i.e. the intermediate zone) at W8 (Fig 2a), in line with our ST data (Fig 1d). We further confirmed early neurogenesis for ExNs (EBF1+), IbNs (PAX2+) and ChNs (ISL1+) by immunohistochemistry (IHC) at W8 (Extended Data Fig 4a-b). A previous study on human developing spinal cord showed that glial cells first appeared at W7-8 ^6^, equivalent to mouse embryonic day (E) 14-16. However, we observed that all glial cell markers were expressed at W5, in which we found that ASCs (*MSX1+GFAP+*) were derived from the dorsal ventricular zone, EPCs (*FOXJ1+RFX4+*) were derived from the ventral ventricular zone, and OPCs (*OLIG1*+*OLIG2*+) were derived from pMN domain (Fig 2a). All these glial cell types showed *MKI67* expression, suggesting that gliogenesis continued during the early stage of first trimester from W5-8 on (Fig 2a, Extended Data Fig 4h). These HybISS data were well correlated with ST and stereoscope data (Extended Data Fig 3), and thus validated the observations by ST. We further performed IHC for these newborn glial cell markers at protein level. We observed IHC signals at W5 for both ASCs (MSX1+GFAP+) and EPCs (RFX4+FOXJ1+) in the dorsal and ventral area of the spinal cord, respectively, in agreement with our HybISS results (Fig 2 b-c). However, using immunofluorescence we did not observe OPC markers at the protein level (PDGFRa+OLIG2+) at W5 (Fig 2d) but found clear double-stained cell profiles at W8 (Extended Data Fig. 4d). Our data suggest that NPCs have were committed to glial fate as early as W5 in the developing human spinal cord.

**Fig. 2:**
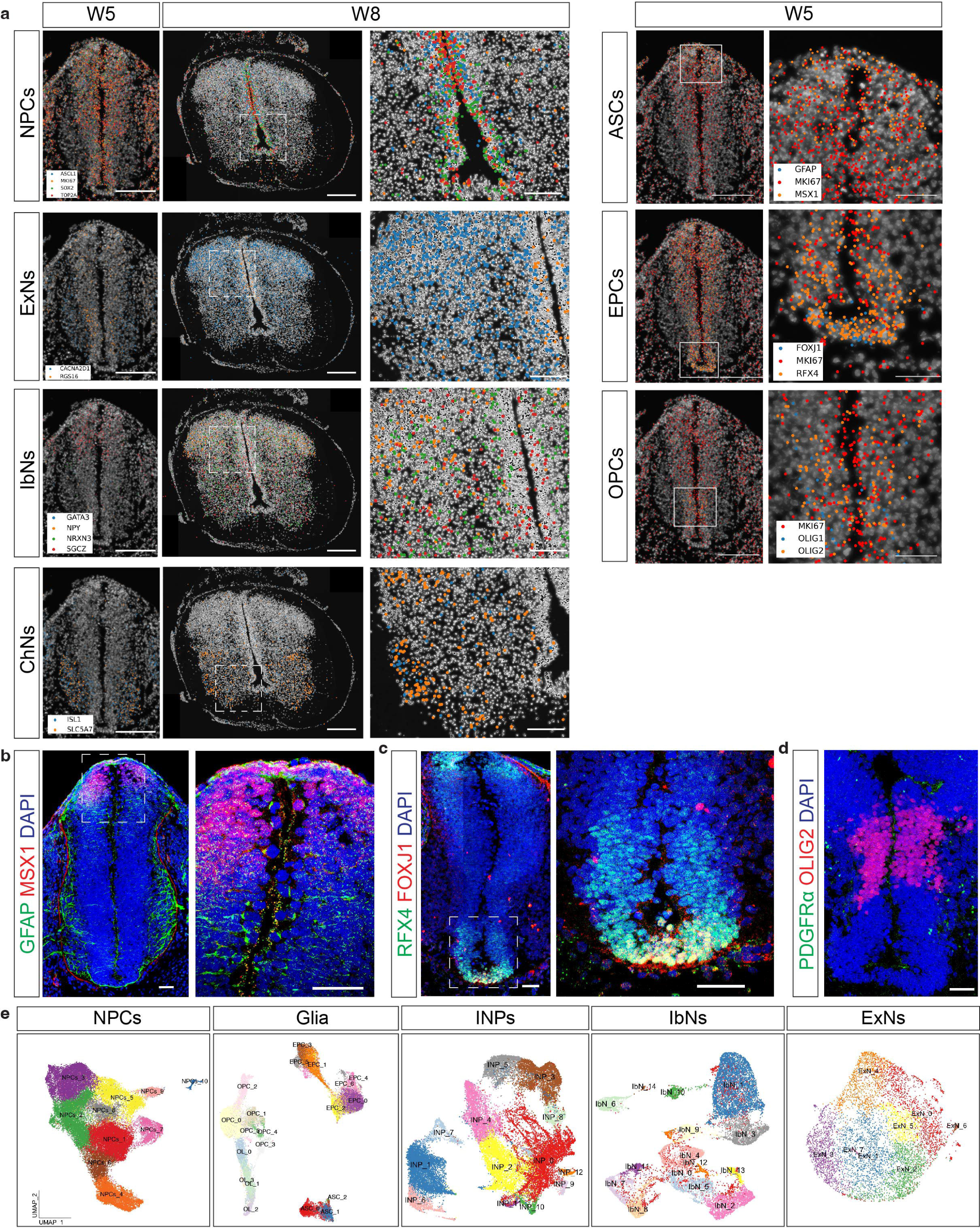
Heterogenous neural cells in the human developing spinal cord. a) Representative images showing validation of newborn neurons and glial cells in the developing human spinal cord by HybISS. Scale bar 200μm. b) Representative confocal images showing immunostaining of newborn astrocytes (b) and ependymal cells (c) at W5, while OPCs are not born at W5 yet (d). Scale bar =200 μm and 50 μm for low and high magnification respectively. Rectangles indicate enlarged areas. e) UMAP illustrating the heterogenous cell types or cell states of different neural cell populations.

To further characterize the heterogenous cell types and cell states during human spinal cord development, we analyzed each major neural cell type and revealed their diversity (Fig 2e). These subpopulations or cell states could be distinguished with single or combinatorial markers (Supplementary Figure 2). We then integrated the scRNA-seq analysis with ST to determine their spatial distribution (Extended Data Fig 5). Some neurons exhibited specific spatial distributions, such as IbNs_2 in the dorsal parts and IbNs_6 in ventral parts. Similarly, the early born glial cells showed specific spatial distributions, with EPCs_0 in the dorsal ventricular zone while EPCs_3 were located in the ventral ventricular zone (Extended Data Fig 5). We further validated the regional distribution of subclusters by HybISS. For instance, IbNs_6 neurons (*TAL2*+) were found in the ventral spinal cord (Extended Data Fig 5; Extended Data Fig 4i), and IbNs_13 neurons in the dorsal-central spinal cord and exhibited *GPC5*, *DTX1* and *ROR1* expression (Extended Data Fig 5, Extended Data Fig 4i). For glial cells, we found that most OPCs were derived from the ventral spinal cord (Fig 2a, Extended Data Fig 5) and were *PDGFRA*+*OLIG2*+, but there were distinct subtypes, such as OPCs_2 (PDGFRA- *OLIG2*+*NKD1*+), and OPCs_3 (*EN2*+) (Extended Data Fig 4i; Supplementary Figure 2). However, many clusters of neuronal and glial cell types did not display regionally specific distributions, suggesting that these subclusters not only include cell subpopulations but also transient cell states during development. Indeed, by performing gene ontology analysis on the differentially expressed genes (DEGs) of different neuronal and glial cell populations, the most common results were associated with “neurodevelopment”, “neurogenesis” and “gliogenesis”. Therefore, we focused on how neurogenesis and gliogenesis are regulated by spatiotemporal gene expression during NPC self-renewal, fate commitment and differentiation in the developing human spinal cord.

### NPCs are committed to neuronal and glial fates during early spinal cord development

In the analysis of neurodevelopment, we first focused on the NPC populations and found 10 different clusters in the scRNA-seq dataset (Fig. 2e). The NPCs clusters could be characterized either by a single marker or by combinatorial markers (Extended Data Fig 6a), and they all expressed neural stem cell and radial glial cell markers at mostly high levels, indicating their stem cell properties (Extended Data Fig. 6b). In contrast to the common view that most NPCs proliferate extensively at this stage ^1^, we found that more than half of the clusters expressed low levels of active cell cycle genes (S or G-to-M phase) (Extended Data Fig. 6c). Spatial distribution analysis showed that hNPCs were mostly located around the ventricular zone but with a relatively smaller area at later timepoints compared with W5 (Extended Data Fig. 6d). From W9 to 12, hNPCs could be observed in the intermediate and marginal zones, indicating migration of newly differentiated neuronal and glial progenitors (Extended Data Fig. 6d). Interestingly, HybISS data showed that the expression of proliferation markers (*MKI67* and *TOP2A*) substantially decreased in hNPCs, and hNPCs_10 had even disappeared from W9 (Extended Data Fig. 6e). In agreement, immunohistochemistry (IHC) showed that many SOX9+ hNPCs did not express KI67 in the W5 human spinal cord, suggesting that a large proportion of hNPCs enter quiescence in early development (Fig. 5a).

To analyze the starting point of differentiation, we used two different methods for trajectory analysis, scVelo^13^ (Extended Data Fig 7a-b) and URD^14^ (Extended Data Fig 7c-e) on the NPC populations. We showed that all NPC populations were highly connected with each other (Extended Data Fig 7a). The proliferative hNPCs (NPCs_5, 7, 9 and 10) changed their fates towards low-proliferating NPC clusters, developed into NPCs_3 and 4, and further into neurons and glia (Extended Data Fig 7b, more details in Fig. 3). By calculating the most significant lineage-associated genes, we found that different genes including TFs were specifically associated with either neuronal or glial lineages (Extended Data Fig 7e), suggesting that most NPCs were genetically regulated for fate commitment into either neurons or glia at W5 in the human developing spinal cord (Extended Data Fig 7c-e). We confirmed these observations by integrating our scRNA-seq dataset with a recent study on early human spinal cord development (from W4 to W7) ^7^ (Extended Data Fig 6f). Consistently, we showed that the most proliferative NPC populations (NPCs_5,7,8 and 9) highly expressed MKI67 and TOP2A, while differentiating NPCs (NPCs_3 and 4) expressed specific markers (*EPHA6* and *SULF2* for NPCs_3, *GADD45G* and *DLL3* for NPCs_4) (Extended Data Fig 6g), in line with the DEG results from our own NPC data (Extended Data Fig 6a). By selecting NPCs from the earliest stages in this integrated dataset (W5 in our data and CS12 from the Rayon2021 data), we analyzed the DEGs by comparing non-proliferative NPCs with proliferative NPCs. Performing GO analysis, we found that the neuronal differentiation and neurogenesis were the top suggested biological processes (Extended data Fig 6h), suggesting that non-proliferative hNPCs were involved in differentiation, in line with our trajectory analysis of NPCs.

**Fig. 3:**
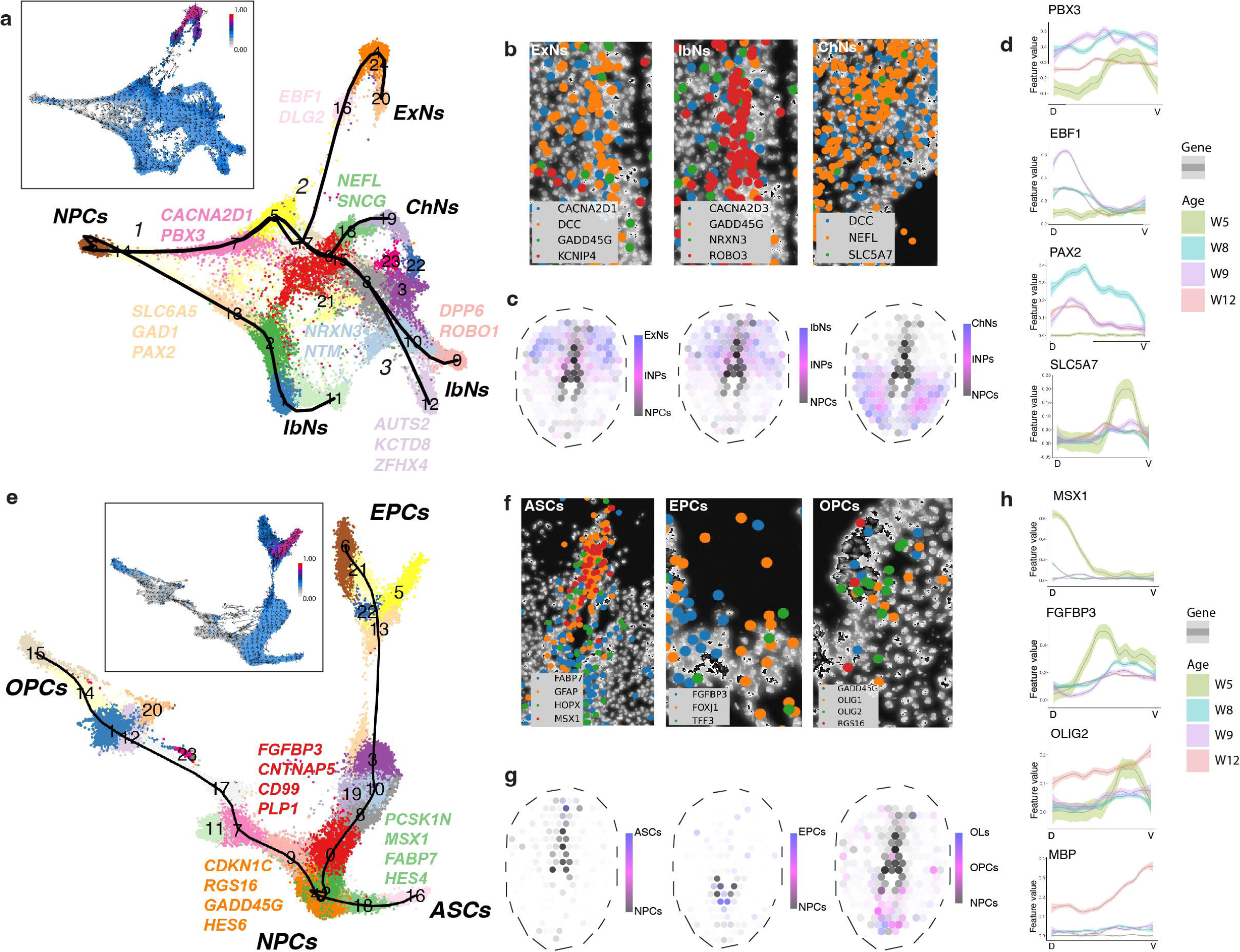
Spatiotemporal regulation of human neurogenesis and gliogenesis. a) UMAP displaying branches from NPCs to different neuronal clusters, confirmed by RNA velocity (left upper panel). Lighter colors - undifferentiated states; darker colors differentiating states. b) HybISS revealing the co-location of NPCs, neuronal markers and lineage-related genes revealed by trajectory analysi. c) Integrated trajectory and ST data revealing neuronal spatial differentiation. d) Spatial quantification of neuronal lineage-associated gene expression along DV axis across ages. e) UMAP indicating branches from NPCs to different glia, confirmed by RNA velocity (upper panel). f) HybISS revealing the co-location of NPCs, glial markers and lineage-related genes. g) Integrated trajectory and ST data revealing glial spatial differentiation. h) Spatial quantification of the expression of glial lineage-associated genes along DV axis across ages.

### Spatiotemporal gene expression regulates neurogenesis and gliogenesis in the human developing spinal cord

To characterize NPC development into different neuronal and glial populations, we selected the clusters related to NPCs and neurons from the scRNA-seq data for three trajectory analysis by Slingshot, RNA velocity and URD ^14–16^ (Fig. 3a and e, Extended Data Fig. 8 a-b). Slingshot analysis revealed that NPCs gave rise to different neurons during neurogenesis (Fig. 3a-b; Supplementary Figure 3a-b), with specific gene expression associated with each branch (Fig. 3a; Supplementary Figure 3c-d). As shown previously in W8-12, ExNs and IbNs mainly occupied the dorsal horns while ChNs were distributed in the ventral area (Extended Data Fig. 3c, Fig. 2a). We validated whether these newborn neurons co-expressed neuronal markers and trajectory-related genes by HybISS and found that at W8 they co-expressed NPC marker genes (*DCC* and *GADD45G*), neuronal-lineage associated genes revealed by trajectory analysis (*CACNA2D1* in ExNs, *NRXN3* in IbNs, *NEFL* in ChNs;) and neuronal markers (*KCNIP4* in ExNs, *ROBO3* in IbNs, *SLC5A7* in ChNs) (enlarged areas shown in Fig. 3b, overview images in Extended Data Fig. 8c). Our results validated that hNPCs co-express specific lineage-associated genes when they are committed into specific neuronal cells.

As neurons have regionally specific distributions in the spinal cord (i.e. sensory neurons in the dorsal horn and motor neurons in the ventral horn), we wondered whether this spatial distribution of different cell types is not only temporal, but also spatially regulated during neurodevelopment. To this end, we spatially delineated neuronal differentiation, by integrating our scRNA-seq trajectory and ST data. We found that hNPC differentiated into INPs first, then into different functional neurons, and the process was specifically regulated to the spatial distribution of different neurons (Fig. 3c). Furthermore, to validate these spatial trajectory calculations, we developed a method and implemented it as an R package that allowed us to spatially quantify gene expression along the DV axis in the ST dataset, independent from the scRNA-seq data. We found that the most significant temporal lineage-associated genes revealed by scRNA-seq, such as *EBF1* (for ExNs), *PAX2* (for IbNs) and *SLC5A7* (for ChNs) exhibited a biased DV expression in ST analysis, which correlated with the terminally differentiated neuron types (Fig. 3d, Extended Data Fig. 8d). *PBX3* was associated with all three neuronal lineages, thus did not exhibit a specific DV pattern from W8 (Fig. 3d). The results were again confirmed by the gene expression pattern displayed with HybISS (Extended Data Fig. 8c). Therefore, the temporal genetic regulation of human neurogenesis revealed by our scRNA-seq data correlates with spatial positioning of neuronal subtypes in the developing spinal cord, revealed by ST and HybISS.

To investigate gliogenesis in the human developing spinal cord, we performed similar analysis as above integrating scRNA-seq and ST. We found that all three glial lineages originated from one common NPC subtype (Fig. 3e, Extended Data Fig. 8a-b) and identified the most significant genes associated with the branches of the trajectories, such as *CNTNAP5* for EPCs, *MSX1* for ASCs and *HES6* for OPCs (Fig. 3e, Extended Data Fig. 8b, Supplementary Figure. 3e-f). Since the newborn glial cells exhibited certain spatial patterns at W5-9 before migration at W12 (Extended Data Fig. 3), we inferred that the temporal lineage-associated genes could also spatially regulate gliogenesis. Indeed, *MSX1* (in ASCs), *FGFBP3* (in EPCs) and *RGS16* (in OPCs) were spatially expressed in the same area as the newborn glial cells i.e. the *GFAP+* ASCs, *FOXJ1*+ EPCs and *OLIG1+* and *OLIG2*+ OPCs (Fig. 3f, Extended Data Fig. 8c). Integrated trajectory and ST data showed that hNPCs differentiated into glial cells in specific spatial patterns – ASCs in the dorsal, EPCs in the central and OPCs and OLs in the ventral spinal cord (Fig. 3g). We further quantified the expression of these top lineage-associated genes along the DV axis and found that *MSX1* was highly expressed in the dorsal spinal cord, *FGFBP3* in central, and *OLIG2* in the mid-ventral domain from W5-8 (Fig. 3h, Extended Data Fig. 8e). The spatial expression pattern of *MSX1* and *FGFBP3* disappeared at W9, suggesting that the patterning of newborn ASCs and EPCs mainly took place before W8. In contrast, *OLIG2* continued to show high ventral expression, while the mature OL-associated gene *MBP* exhibited strong ventral expression at W12, which correlated with the appearance of newborn mature OLs in the ventral spinal cord at W12 (Extended Data Fig. 3). These data were validated by both ST (Extended Data Fig 8e) and HybISS (Extended Data Fig. 8c).

To further analyze the active TFs that regulate cell fate commitment, we performed regulon analysis by SCENIC ^17^ in the scRNA-seq dataset. The analysis revealed the top regulons for human spinal cord development as well as the gene expression of the top TFs (Extended Data Fig. 9a-b), and we found that the top regulons were active in specific lineages during neurogenesis and gliogenesis (Extended Data Fig. 9d). Most of the regulons for glial cells had been active since W5 (Supplementary Figure 5a), indicating that NPCs were committed not only to neuronal but also to glial fates at this early stage. This is also in line with gene expression and validation by HybISS and IHC data above for early glial cells at W5 (Fig 2 a-d, Extended Data Fig. 4h). Altogether, our analysis showed that the fate commitment of hNPCs is spatiotemporally regulated by specific gene sets in the developing human spinal cord.

### The spatiotemporal genetic regulatory networks of human spinal cord development

During neurodevelopment, hNPC differentiation follows a spatiotemporal pattern of cell fate commitment, and is defined in specific progenitor and neuron domains along the DV axis ^1, 18^. To better understand the regulatory controls (e.g., expression of TFs, morphogens, signaling pathways, cell-cell interactions etc.), we first surveyed the most well-known signaling pathways that have an impact on neural patterning, and found that the gene modules of WNT, NOTCH and SHH signaling were expressed by most cell types (Fig. 4a) but overall decreased overtime (Fig. 4b, Supplementary Figure 4). IHC showed that Active-β-Catenin (ABC) and SHH pathway molecules (SHH, GLI1 and GLI3) at W5 were expressed in the roof plate and floor plate respectively (Fig. 4c, Extended Data Fig 4e-f), but the expression decreased dramatically after W8 (Fig. 4c). However, the NOTCH target HES1 showed overall high expression level throughout the ventricle layer, without much DV biased expression (Extended Data Fig 4g). Under the gradients of morphogens such as SHH and WNT, genes associated with neurogenesis and gliogenesis (revealed by trajectory analysis) exhibited spatially specific expression pattern at W5 and W8, which coincides with the spatial positions of their related differentiated cell types, shown by HybISS (Fig. 4d). As variable genes used for analysis in the scRNA-seq are dominated by differentiation during development, it was challenging to spatially investigate and quantify the expression of neural patterning genes ^7, 19^. We used ST to directly measure the expression of neural patterning genes, and created a detailed spatial gene expression panel that indicates the DV patterning in the developing human spinal cord at early stage (W5-8) (Fig 4e, Extended Data Fig. 10). Further, we quantified the expression of these neural patterning genes, and showed that their spatially unique expression was also restricted to certain developmental stages (i.e. progenitor patterning genes showing DV biased expression at W5 and neuronal patterning at W5-8) (Fig. 4e).

**Fig. 4:**
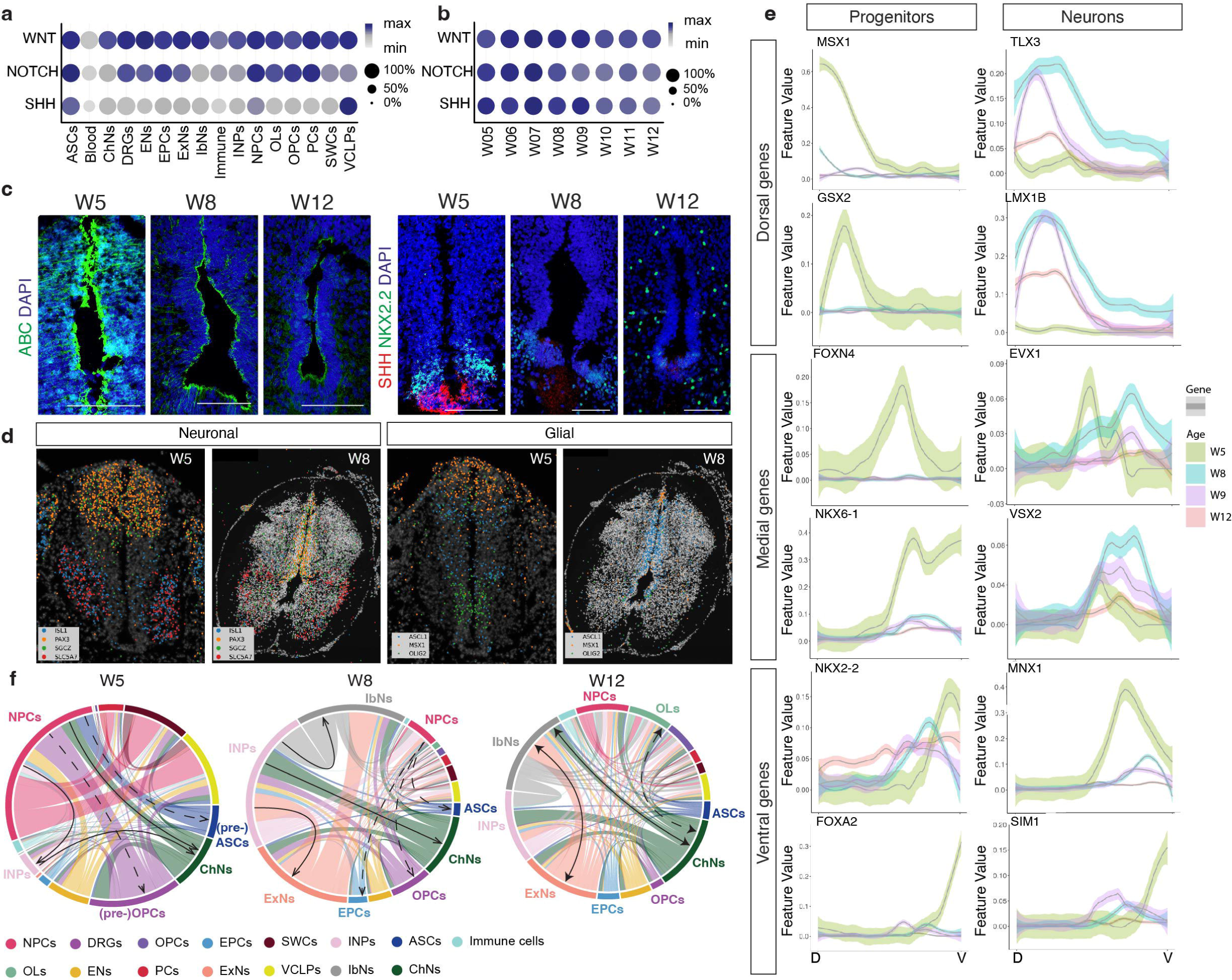
The regulatory networks of human spinal cord development. a-b) Dot plots illustrating the expression of the three major signaling pathways involved in spinal cord development in different cell types (a) and their decreased expression during development (b). Max = highest expression of the given gene or module; Min = 0. c) Representative confocal images of immunostained active-β-catenin (ABC) and SHH during human spinal cord development from W5-12. d) HybISS revealing neuronal and glial progenitor patterning. e) Examples of spatial quantification of neural patterning genes along DV axis across ages. Scale bar 100 μm. f) Circos plots displaying colocalization and major connections of different cell types during development. Solid lines: neurons; dashed lines: glia.

Since multiple cells contribute to each spatial capture location in ST, we performed a co- localization analysis using the proportion estimates obtained from *stereoscope*. This allowed us to visualize the initiation of cell fate transition locally before migration starts. We found that the ratio of NPCs decreased dramatically during development (Fig. 4f). At W5, the major connections between NPCs to neurons and pre-glial cells represent initial differentiation. At W8, the shared locations between INPs and different neurons suggested ongoing local neurogenesis preceding neuronal migration. In contrast, the connections between NPCs and glial cells were weak from W8, suggesting that soon after glial fate commitment migration of the immature glial cells begins, during which further differentiation takes place (Fig. 4f). At W12, the weak connections between NPCs and others, and strong connections among neurons and glia suggested that the major events had shifted from differentiation to the formation of neural circuits. In addition, TFs and cell-cell interaction analysis revealed other regulatory networks such as top TFs in each cell type and cell-cell integrations via the most significant ligand-receptor interactions (Supplementary Figure 4). This network analysis was in line with our *in situ* data showing co-locolization of NPC and neural cell markers at early developmental stages (Fig. 3b and f, Extended Data Fig. 8c).

### Neurodevelopment involves species-specific events

Most NPCs are believed to proliferate extensively before gliogenesis starts^1^, we found that more than half of the clusters expressed low levels of active cell cycle genes (S or G-to-M phase) (Extended Data Fig. 6c). To address whether low proliferation is a specific phenotype in humans, we integrated our scRNA-seq datasets with two mouse spinal cord development datasets ^19, 20^ for normalized gene expression comparison (Fig 5a). In contrast to the majority of hNPCs that had low expression of proliferation markers *MKI67* and *TOP2A* from W5-7, mouse (m)NPCs were highly proliferative at least up to embryonic day (E) 13.5 (equivalent to human W7) (Fig. 5b). One main mechanism that drives the quiescence of NPCs is the molecule *LRIG1* ^21^, which showed much higher expression during embryonic and fetal stages of the human spinal cord compared to mouse, in which high expression took place postnatally (Fig. 5b). In agreement, IHC showed that many SOX9+ hNPCs were not expressing KI67 in the W5 human spinal cord, differently from mouse E10.5 spinal cord containing mostly Sox9+Ki67+ cells, confirming that most hNPCs in contrast to mNPCs entered quiescence during early development (Fig. 5c).

**Fig. 5:**
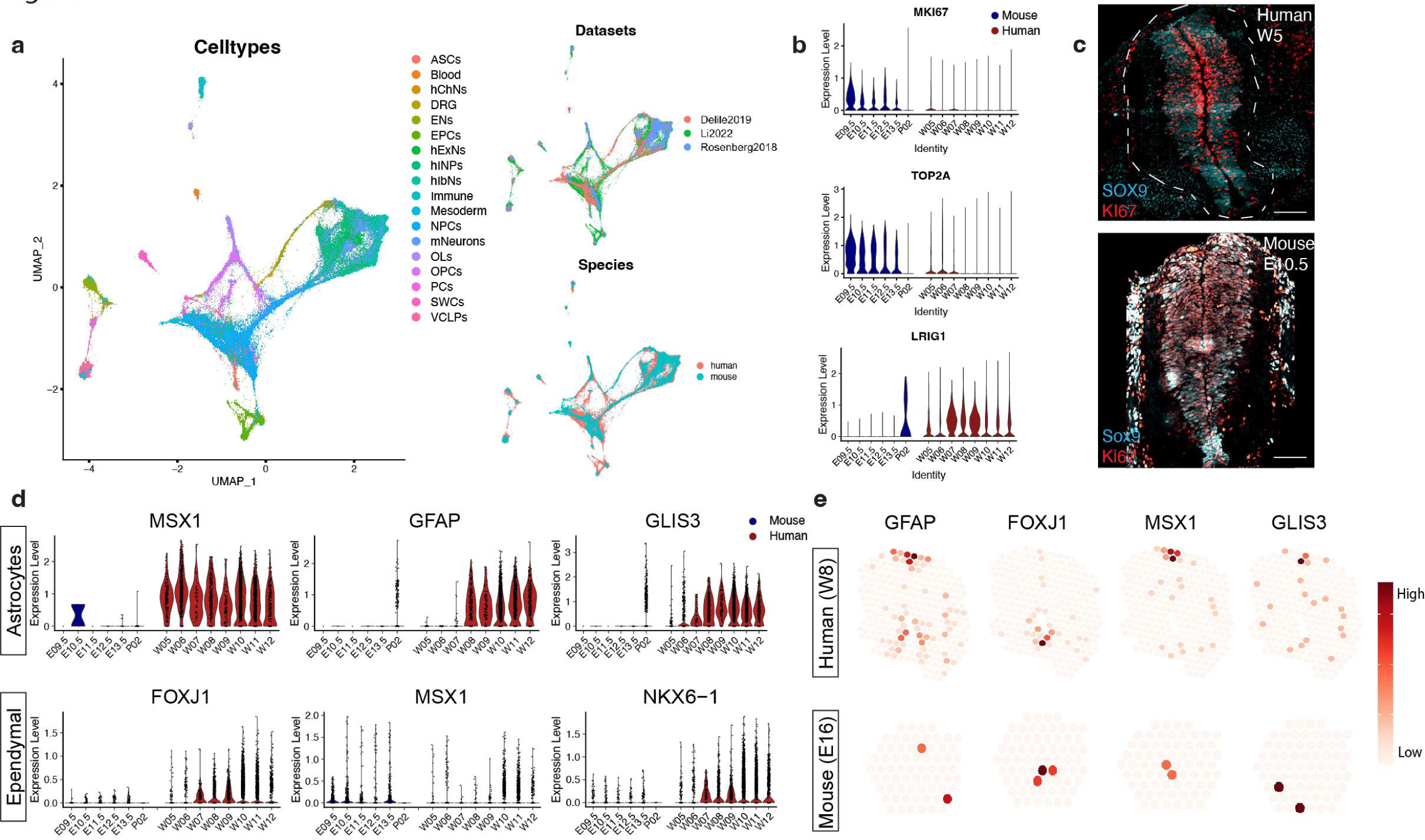
Species-specific events during neurodevelopment. a) UMAP illustrating integrated scRNA-seq datasets of human and mouse spinal cord development. Li2022: dataset in this study. Other two datasets: publicly available mouse development datasets. b) Violin plots displaying normalized gene expression of proliferation markers *MKI67* and *TOP2A*, as well as stem cell quiescence regulator LRIG1 during mouse and human spinal cord development. c) Representative confocal images illustrating proliferative human and mouse NPCs at early stage. Scale bar = 100 μm. d) Violin plots displaying species difference gene expression of gliogenesis regulators. e) ST plots displaying differences of spatial gene expression of gliogenesis regulators in the developing spinal cord between human and mouse.

Recent studies with scRNA-seq reveals that ASCs and OPCs are derived as early as gestational week 8 (equivalent to post-conception week 6-7 in this study) in the developing human spinal cord ^6^. By using HybISS to validate lineage associated genes revealed by scRNA-seq, we found these lineage-associated genes had been expressed in glial cells (*MSX1* in ASCs, *OLIG1* and *OLIG2* in OPCs, and *FOXJ1* in EPCs) as early as W5, equivalent to E11 in mouse development. It is known that the first mouse ASCs, OPCs and EPCs appear at E16.5, E12.5 and E15.5 respectively ^22–25^, suggesting that hNPCs might have been committed to gliogenesis at earlier stage of human neurodevelopment. Interestingly, *Msx1* has been shown to be the key regulator for EPC differentiation during mouse spinal cord development ^26^, and we found that *MSX1* is both a cell marker and a lineage-associated gene for human ASCs (Fig 1c, Fig 2a, Fig 3e-f). By comparing gene expression in both ASCs and EPCs in the human-mouse integrated scRNA-seq dataset, we found that *MSX1* was indeed highly expressed in *GFAP* expressing ASCs in human spinal cord, but had low expression levels in mouse ASCs during development (Fig 5d). In contrast, *MSX1* and *FOXJ1* were expressed in both mouse and human EPCs, suggesting that *MSX1* has dual roles in regulating cell fate commitment of human ASC and EPC, but only regulates EPCs in mice. We validated these results by performing ST at comparable timepoints of human (W8) and mouse (E16) spinal cord sections. We found that the expression of *MSX1* in humans was mainly located in the dorsal ventricular zone, and correlated with the marker gene expression *GFAP* in ASCs and *FOXJ1* in EPCs (Fig 5e). However, in mouse sections, *Msx1* was found to be expressed in the same area of *Foxj1*+ cells, but not in *Gfap*+ area, in line with the previous study ^26^ showing *Msx1* is expressed in mouse ependymal cells during development (Fig 5e).

To further compare the similarities and differences between mouse and human development, we compared the most significant TF activities in human and mouse developing spinal cords by regulon analysis (Extended Data Fig. 9 a and c). We found that some specific regulons associated with certain human cell types (Extended Data Fig. 9a) also showed gene expression in these cell types, but not in mice. For instance, *GLIS3* was highly associated with the fate commitment of ASCs in human but little in mice, while NKX6-1 was expressed in human EPCs regulating their development but not expressed in mouse EPCs (Fig 5d). We further confirmed these findings by ST, and showed that *GLIS3* was associated with *GFAP*+ area in human but not mouse developing spinal cord (Fig 5e). Altogether, our data suggest that despite the conserved mechanisms, there are fundamental differences of spatiotemporal gene expression between mouse and human spinal cord development.

### Fetal human spinal cord and relation to ependymomas

Ependymomas are highly aggressive CNS tumors with high recurrence rate ^27, 28^. Furthermore, childhood ependymomas show much higher rates of anaplastic histology ^29^, suggesting a larger CSC population in the tumor. It has been suggested that the development of pediatric ependymomas recapitulates neurodevelopment, but the previous studies on pediatric ependymomas progression lacked proper normal human neurodevelopment datasets as control^27, 28^. To demonstrate the potential of the human developmental atlas to reveal novel biomarkers for disease diagnosis and potential treatment, we used our data to gain insight into the molecular signature and differentiation of drug-resistant cancer stem cells (CSCs) in pediatric ependymomas. We first obtained genes related to spinal cord tumor (HP:0010302) from the Human Phenotype Ontology (HPO) database and plotted the module on ST data. We observed broad but no regionally specific gene module expression of spinal cord tumor in all ST sections (Fig. 6a), suggesting that many cell types in normal human developing spinal cord are highly similar to spinal cord tumors. We integrated our scRNA-seq data with human pediatric ependymomas ^28^, unbiasedly transferred cell types from normal developmental data to ependymomas, and showed that many cell types in normal tissue and in tumors were well integrated (Fig. 6b, Supplementary Figure 6a). Despite the overlapping clusters of neural-cell-like ependymoma with normal cells (Fig. 6c and f), most of the normal neuronal markers were predominantly expressed in the normal neurons (Fig. 6d) while the normal glial markers were similar between normal and tumor cells (Fig. 6g). The non-overlapping area probably suggests that certain cell populations are unique to the respective condition (normal vs tumor). In the field of cancer diagnosis and treatment, it is usually challenging to separate tumor and normal cells to identify cancer-specific biomarkers. Therefore, we focused on comparing the overlapping clusters between normal and tumor data, and identified tumor-specific genes such as *CASC15* and microRNA *MIR99AHG* in neuron-like ependymomas, and *RPS14* and *RPS8* in glia-like ependymomas (Fig. 6e, h). Moreover, many CSCs overlapped with normal NPCs (Fig. 6g) and shared the expression of the classical NPC markers *SOX2* and *VIM* (Fig. 6j). After identifying the putative CSC markers-associated clusters and proliferative clusters (cluster 3, 6, 7) (Supplementary Figure 6c, Fig 6j), we uncovered the CSC specific markers *FTX* and *MIR99AHG*, which were not expressed in the normal hNPCs and thus could be novel therapeutic targets for the ependymoma CSCs (Fig. 6i-k). We also plotted these CSC-specific genes (e.g. *FTX* and *MIR99AHG)* in the ST dataset from normal human spinal cord sections as independent validation, and did not find any expression in the sections (data not shown), confirming that these genes are tumor-specific.

**Fig. 6:**
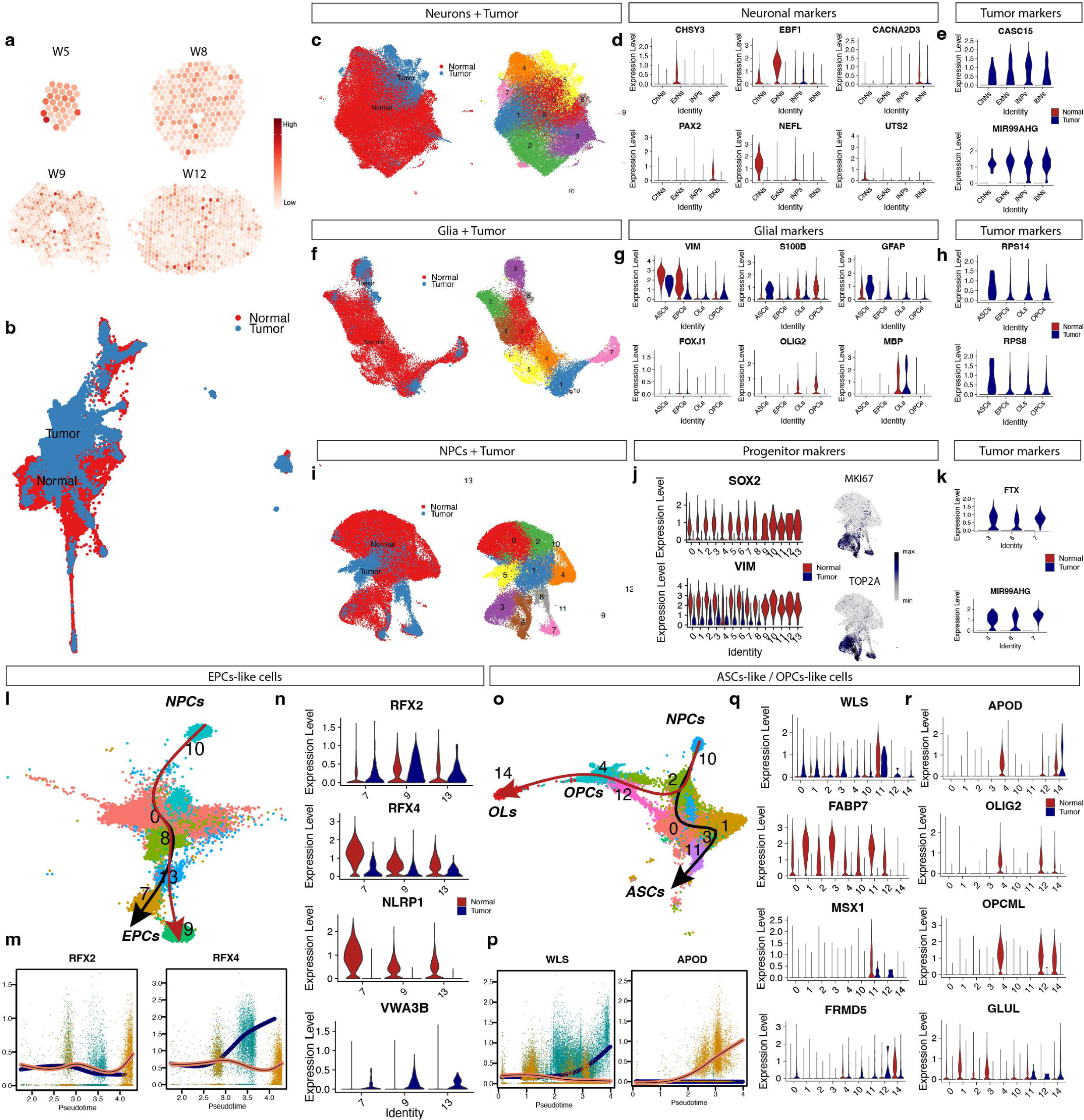
Fetal human spinal cord and relation to ependymomas. a) ST plots displaying spinal cord tumor gene module expression. b) UMAP displaying integrated normal human spinal cord and ependymomas scRNA-seq datasets. c-e) Clusters of neuronal populations shared between conditions (c) and the expression of normal neuronal markers (d) and tumor specific markers (e). f-h) Clusters of glial populations shared between conditions (F) and the expression of normal glial markers (g) and tumor specific markers (h). i-k) Clusters of progenitor populations shared between conditions (i) and the expression of normal stem cell markers (j) and tumor specific markers (k). l-n) Trajectory analysis of EPCs- like cells (l) and lineage-associated gene expression along pseudotime (m) or among branch-related clusters (n). o-r) Trajectory analysis of ASCs-like and OPCs/OLs-like cells (o) and lineage-associated gene expression along pseudotime (p) or among branch-related clusters (q- r).

A previous study suggested that ependymoma-derived CSCs mimics neurodevelopment ^28^. To further investigate the molecular differences between normal and cancer progenitors during differentiation, our trajectory analysis showed that the EPC-related TFs *RFX2* and *RFX4* were highly associated with two lineages of ependymal cell differentiation (Fig. 6l-m) and were associated with both normal and tumor EPC differentiation. By screening the top lineage-associated genes (Supplementary Figure 6b), we found that *NLRP1* and *VWA3B* were specifically associated with normal ependymal and ependymal-like cancer cell differentiation, respectively (Fig. 6n). Similarly, we found that *WLS* and *APOD* were highly associated with the differentiation of ASCs (or ASC-like tumor) and OPCs (or OPC-like tumor) respectively (Fig. 6 o-p). However, *FABP7* and *MSX1* were mainly associated with normal ASCs and *OLIG2* and *OPCML* were mainly associated with normal OPCs and OLs (Fig.6 q-r, Supplementary Figure 6b). In contrast, *FRMD5* and *GLUL* were mainly associated with ASCs- like and OPCs/OLs-like ependymomas (Fig. 6 q-r, Supplementary Figure 6b) respectively. Altogether, with our human spinal cord developmental atlas, we provide new insights of normal human spinal cord development and potential diagnostic or therapeutic strategies in human CNS tumors.

### An integrated atlas of spinal cord cell types across rodents and humans

To compare cell type differences across species, timepoints, and technologies, we performed stepwise data integration to integrate our human scRNA-seq data with publicly available scRNA-seq datasets of spinal cord samples, including human development ^6, 7^, mouse embryonic development ^19^, mouse postnatal development ^20^, mouse adulthood ^30–34^ and datasets suggested in a meta-analysis of mouse spinal cord atlases ^35^. We created an integrated cell atlas with these 1.8 million cells (Fig 7a-c). We compared our cell type annotation results to the original annotation from several datasets and found high correlations (Supplementary Figure 7). We then performed label transferred from our annotated cell types to the integrated dataset (Fig 7a). Using the cell proportion from our dataset as a baseline for dataset comparison, we found that our dataset (Li2022) shared high similarity with other comparable datasets of mouse and human development, as shown in Zhang2021, Delile2019 and Rayon2021 (Fig 7d). Notably, the Zhang2021 dataset includes some samples from second trimester of human spinal cord development, but not much difference in OPCs and EPCs, suggesting a continuation of glial cell differentiation but probably few newborn glial cell progenitors during the second trimester in the developing human spinal cord. This large integrated dataset is now also available together with the interactive map of our multi-omics data (https://tissuumaps.scilifelab.se/web/HDCA/SpinalCord2022/index.html).

**Fig. 7:**
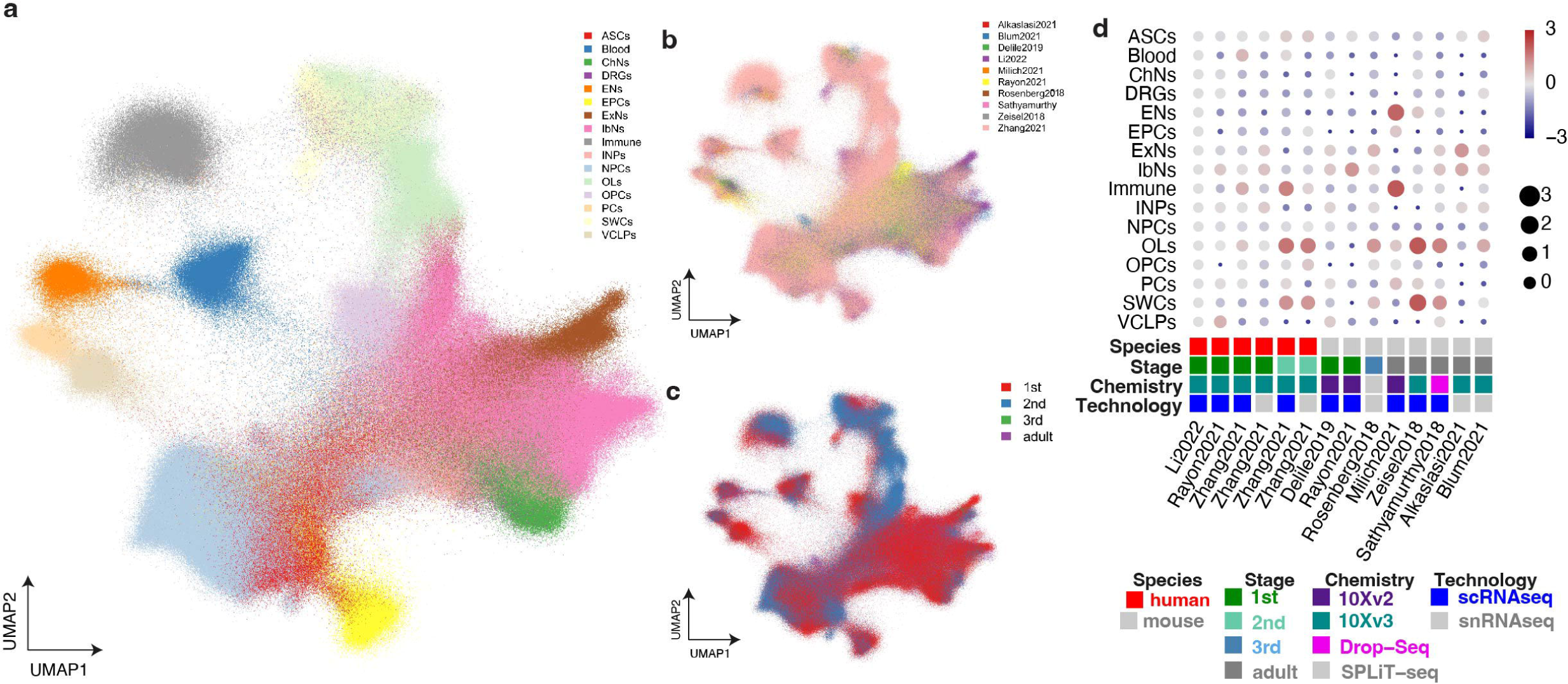
An integrated atlas of spinal cord cell types across rodents and humans. a-c) UMAP illustrating the integrated spinal cord scRNA-seq dataset with cell types (a), and across datasets (b) and developmental trimesters (c). d) Dot plot illustrating cell proportions across different species, developmental stages, cell capturing chemistry and technologies.

## Discussion

In this study, by using multi-omics and data integration to study the developing human spinal cord, we have: i) created a developmental cell atlas of the human spinal cord throughout the first trimester of development, ii) revealed spatiotemporal regulation of human spinal cord neurogenesis and gliogenesis, iii) presented major differences of cell and molecular regulation between rodent and human spinal cord development, and iv) discovered unique markers and regulation of CSC differentiation in human ependymomas.

The dynamics and molecular regulation of the human spinal cord development are still understudied. While two recent studies explored the developing human spinal cord by scRNA-seq and showed neural patterning and neurogenesis in identified clusters, they did not elucidate how NPCs are committed to multiple neural cell lineages or how the spatiotemporal gene expression is involved in neurogenesis and gliogenesis ^6, 7^. In this study, we acquired human prenatal spinal cords over the first trimester for scRNA-seq and spatial techniques, integrated the multi-omics datasets and validated the results, which gave new insights into the spatiotemporal gene expression of the developing human spinal cord.

NPCs are believed to proliferate vividly during fetal development ^1^. However, we found that many hNPCs throughout the ventricular zone did not proliferate even at the early embryonic stage. The proliferative NPCs lose their proliferation during the first trimester in humans, much earlier than rodents. The loss of active NPCs after fetal development limits regeneration in the mammalian adult spinal cord, for example after SCI ^24^. The loss of active NPCs during first trimester development in humans partly explains the extremely low regenerative potential in human spinal cord. Interestingly, our results are in agreement with a previous study that the stem cell quiescence-related gene *LRIG1* ^21^ is one of the most significant gene in the non-proliferative hNPCs, which could be targeted for reactivating hNPCs.

Since most lineage tracing techniques cannot be applied in humans, it is unclear how neurogenesis and gliogenesis in human spinal cord are regulated in a spatiotemporal manner. With integration of multi-omics data, we highlighted some unique developmental events in the human developing spinal cord. First, by analyzing marker gene expression and active TFs, validated by HybISS and IHC, we found that hNPCs were committed to glial fates as early as W5 while previous studies on active regulons and marker expression showed that this occurred at W8-10^1, 24, 36^. Thus, our data pushes human gliogenesis to an earlier stage of neurodevelopment. Second, while rodent astrocytes migrate horizontally during development to the mantle zone and the future lateral white matter ^36^, we showed that human astrocytes were first restricted in the dorsal region of the spinal cord. In addition, this process in humans was spatiotemporally regulated by *MSX1*, a TF shown to specifically regulate ependymal cell development in rodents ^26^. Third, we conclude that human EPCs exhibit a longer developmental period than expected. Mouse spinal cord EPCs are derived from mid-late fetal stage (E15.5) and are fully developed within one week *in vivo* ^24^. However, while we found that human EPCs are derived from W5, a subpopulation that is located in the dorsal central canal of the human adult spinal cord ^37^ was still missing at W12, suggesting a second wave of gliogenesis during the second trimester. However, by comparing our data with data from Zhang et al., 2021^6^ that includes second trimester samples, we did not observe a significant increase in the proportion of EPCs (Fig. 7d). This could be due to a significant increase of other cell types during second trimester, or technical issues of capturing EPCs during single cell collection from the second trimester samples in other datasets. Future studies involving scRNA-seq and spatial techniques is needed to fully describe gliogenesis in the human spinal cord. Importantly, by comparing genetic regulation of human and mouse neurogenesis and gliogenesis, we found a number of regulons and expressed genes that are only present in human spinal cord development and not in mice, suggesting that neurodevelopment is regulated differently between species. Notably, while most studies on neurogenesis and gliogenesis have focused only on the temporal gene expression as molecular mechanisms, we developed a method to demonstrate that neural patterning and positioning of neural cells are the results of the spatially biased expression in addition to temporal gene expression.

Finally, we applied our developmental atlas of the human spinal cord to investigate gene expression in childhood spinal ependymomas. Pediatric ependymomas are CNS tumors with high recurrence rates, probably due to the proliferation of drug-resistant CSCs ^28^. In the field of drug discovery and development, it is challenging to find cancer-specific markers to selectively target cancer cells. We integrated our human spinal cord atlas with human pediatric ependymomas data, and displayed the most significant differences of gene expression between cancer and normal stem cells. Although CSCs and NPCs share similar lineage-associated genes during differentiation, we could identify unique genes associated with normal NPC and CSC differentiation respectively. Therefore, our results revealed molecular signatures of CSCs, and their potential regulators at different stages during differentiation, which gives new insights into specific targets for ependymoma treatments.

In conclusion, we provide a comprehensive analysis of the human first trimester spinal cord during a critical phase of cellular specification and differentiation. While we confirm that humans and rodents share multiple similar cellular and molecular mechanisms during neurodevelopment, we discovered unexpected unique developmental events in the human spinal cord. Our database will not only serve as developmental cell atlas resource, but also provide important information for research on human neurodevelopmental disorders as well as regenerative strategies and cancer treatments.

## Methods

### Human prenatal tissue

16 samples of human prenatal spinal cord tissue were used in the study (13 for scRNA-seq and 6 for ST, HybISS and IHC) representing post-conception weeks (W) 5, 6, 7, 8, 9, 10, 11, 12. W5-8 is referred to as early stages (embryonic) while W9-12 is referred to as later first trimester stages (fetal) in the present study. Post-conception age was determined by information from the clinical ultrasound, time for last menstrual period, and by identifying age-dependent anatomical landmarks with true crown-rump-length (CRL), taking into account that post-conception age and clinical age differs by 1.5–2 weeks. The prenatal specimens were retrieved from elective medical abortions at the Department of Gynecology, Danderyd Hospital and Karolinska Huddinge Hospital after oral and written informed consent by the patient. All patients were at least 18 years of age and Swedish-speaking. The clinical staff that informed the patients and performed the abortions did not in any other way participate in this research. The specimens were transported immediately from the clinic to the dissection laboratory. spinal cord tissue was rapidly dissected in 4°C saline (Fresenius Kabi, B306443/01) under sterile conditions within one-two hours after the abortion. Specific information can be found in Supplementary Table 1. The use of prenatal tissue for this study was approved by the Swedish Ethical Review Authority and the National Board of Health and Welfare. All procedures met the ethical stipulations of the WMA Medical Ethics Manual and the Declaration of Helsinki, and all experiments were performed in accordance with relevant guidelines and regulations.

### Preparation of human prenatal spinal cord for multi-omics

#### scRNA-seq experiment

The W5-W7 spinal cord tissues were used as one piece while W8-12 spinal cords were divided into three pieces (cervical, thoracic, and lumbar regions) before dissociation. The dorsal root ganglia were removed by cutting the roots. Each piece of tissue was minced into smaller pieces using sterile blades and scissors. Artificial cerebrospinal (aCSF) was prepared as previously described ^16^, with modification for Ca_2_Cl_2_ (1 mM) and MgCl_2_ (2 mM). The aCSF was oxygenated with 95% O_2_:5% CO_2_ for 20 min at 4°C. The samples were then digested at 37°C in aCSF. Papain solution (Worthington Biochemical; cat. no. LK003178; 20 U/ml in CSF) and DNase I (Worthington Biochemical; cat. no. LK003172; 1mg/ml) were added to the aCSF to dissociate the tissue. Incubation time was adjusted based on developmental stage, ranging from 15-25 minutes. The spinal cords were subsequently dissociated manually with fire-polished glass pipettes. When most of the tissue was dissociated into single cells, the solution was filtered using a 30µm cell strainer (CellTrics, Sysmex, 04-0042-2316) and collected in a 15-ml Falcon tube. The digestion solution was diluted with 7.5 ml of aCSF and centrifuged at 300g for 5 min at 4°C. The pellets were resuspended in aCSF and transferred to Eppendorf tubes pre-coated with 30% BSA (Sigma-Aldrich, 9048-46-8). After cell counting, the single cell solution was diluted to a concentration of 800-1200 cells/µl and kept on ice for immediate chip loading.

#### ST and ISS experiments

Human spinal cord tissues at W5, 8, 9 and 12 were embedded in Tissue-Tek (OCT) and snap-frozen using an isopentane/dry ice slurry. W8-12 samples were first divided into cervical, thoracic and lumbar. To enable spatial protein and gene expression analyses, the spinal cords were cryosectioned at 16 μm thickness and alternatingly placed on Superfrost microscope glass slides (Thermo Fisher Scientific) and Visium spatial gene expression slides (10x Genomics), after which they were stored at -80°C for no more than XXX days before being used.

#### Immunohistochemistry

Immunohistochemistry was performed as previously described ^24^. Briefly, tissue sections were rehydrated by 1X PBS for 5 min, then primary antibodies diluted in blocking solution (10% normal donkey serum in PBS) were applied to the sections and incubated at room temperature overnight. Secondary antibodies were applied to sections after 2 times wash with 1X PBS. DAPI was applied on sections for 1 min. Sections were mounted after washing and ready for confocal imaging by Zeiss LSM 700.

### Library preparation and sequencing

#### scRNA-seq experiments

Droplet-based single-cell RNA sequencing was performed using the 10x Genomics Chromium Single Cell Kit v3. Single-cell suspensions concentrated at 800-1200 cells/ml were mixed with master mix and nuclease-free water according to the Chromium manual, targeting 5000 cells per reaction. The library preparation and sequencing were done according to the Chromium v3 standard protocol. Sequencing was performed using the Illumina NovaSeq 6000 with an average 152,486 reads/cell.

#### ST experiments

Spatial Gene Expression libraries were generated using Visium Spatial Gene Expression kit from 10x Genomics (https://support.10xgenomics.com/spatial-gene-expression). Each Visium barcoded glass slide contained 4 capture areas, each with ∼5000 spots, and every spot contained probes consisting of a spatial barcode, a unique molecular identifier (UMI) and a poly-dT-VN sequence allowing for mRNA capture. The diameter of each spot was 55 µm and the center-to-center distance between the adjacent spots was 100 µm. Several sections of the same post-conceptional week were placed in each capture area, the number depending on the size of each section. Sections were fixed for 30 min in methanol, stained with hematoxylin and eosin and imaged using Metafer Slide Scanning system (Metasystem, Altlussheim, Germany). Optimal permeabilization time for spinal cord sections was determined to be 20 min using 10x Genomics Visium Tissue Optimization Kit. In total, Visium Spatial Gene Expression libraries from 76 spinal cord sections were prepared by following the manufacturer’s protocol. Libraries were sequenced using Illumina platform (NovaSeq6000, NextSeq2000). The number of cycles for read 1 was 28 bp and 120 bp for read 2.

#### HybISS

HybISS was performed as reported by Gyllborg et al. ^5^. Briefly, after fixation, sections were permeabilized with 0.1 M HCl and washed with PBS. SecureSeal™ Hybridization Chambers (Grace Bio-Labs) were applied around tissue sections and cDNA synthesized by reverse transcribing overnight with reverse transcriptase (BLIRT), RNase inhibitor, and priming with random decamers. The next day, sections were post-fixed before PLP hybridization and ligation at a final concentration of 10 nM/PLP, with Tth Ligase and RNaseH (BLIRT). This was performed at 37°C for 30 min and then moved to 45°C for 1.5 h. Sections were washed with PBS and RCA was performed with phi29 polymerase (Monserate) and Exonuclease I (Thermo Scientific) overnight at 30°C. SecureSeal chambers were then removed. Bridge-probes (10 nM) were hybridized at RT for 1 h in hybridization buffer (2XSSC, 20% formamide). This was followed by hybridization of readout detection probes (100 nM) and DAPI (Biotium) in hybridization buffer for 1 h at RT. Sections were washed with PBS and mounted with SlowFade Gold Antifade Mountant (Thermo Fisher Scientific). After each imaging round, coverslips were removed and sections washed 5 times with 2XSSC and then bridge-probe/detection oligos were stripped with 65% formamide and 2XSSC for 30 min at 30°C. This was followed by 5 washes with 2XSSC. Then the process was repeated for the next cycle of bridge-probes hybridization.

For subtype / cell state markers, Kits from 10x Genomics were provided along with an accompanying protocol (High Sensitivity kit). In summary, the tissue was fixed, and then the direct RNA probe mixture was added (incubated overnight at 37°C). The section was subsequently washed and ligation mix was added (incubated at 37 °C for 2 h). Following washing, rolling circle amplification was performed at 30°C overnight. Lastly, rounds of labeling and stripping were done for detection.

Imaging was performed with a Leica DMi8 epifluorescence microscope equipped with an LED light source (Lumencor® SPECTRA X), sCMOS camera (Leica DFC9000GTC), and 20× objective (HC PL APO, 0.80). Each field-of-view (FOV) was imaged with 24 z-stack planes with 0.5 μm spacing and 10% overlap between FOVs.

### Sequence alignment and annotation

#### scRNA-seq experiments

Single-cell sequencing data were processed using the CellRanger pipeline (version 3.0.2; 10x Genomics). Reads were mapped against the human genome (ENSEMBL genome assembly, release 93) and annotated with GENCODE gene annotations for the GRCh38-3.0.0 genome assembly (GENCODE release 32). Using the BAM files from CellRanger, molecules were mapped into spliced and unspliced transcripts using velocyto (0.17.17) into which loom files were generated for each sample.

#### ST experiments

Sequenced Spatial Transcriptomics libraries were processed using the Space Ranger v1.0.0 pipeline(10X Genomics). Reads were aligned to the human reference genome (ENSEMBL genome assembly, release 93) and annotated using GRCh38-3.0.0 to obtain expression matrixes.

### Data quality and filtering

#### scRNA-seq experiments

The single-cell count matrix was first enriched for protein-coding RNA and lincRNA gene types. Cells with fewer than 500 genes, and genes expressed in fewer than 15 cells were excluded from the analysis. Cells with over 25% mitochondrial gene expression were also excluded.

#### ST experiments

In total, 76 tissue sections were analyzed, resulting in 20835 spots used for data analysis. The count matrix was enriched for protein-coding and lincRNA genes. Count matrix was filtered for all hemoglobin related genes, *MALAT1*, mitochondrial and ribosomal protein coding genes. Spots with fewer than 500 genes and genes expressed in fewer than 5 spots were excluded from analysis of the three post-conception time points.

### Data analysis

#### Analysis for scRNA-seq and ST data

Normalization, dimensionality reduction, and clustering of scRNA-seq data were performed using the Seurat package (Seurat v4.0.1) ^38^, and the top 6000 genes with high dispersion were selected using the FindVariableGenes function. Cell cycle activity, number of genes, and mitochondrial content across the data were regressed out using the ScaleData function. Principal component analysis (PCA) was performed on the 50 most significant components as determined by the PCElbowPlot function, showing the standard deviation of the principal components. Cells in different cycling stages were identified by gene sets called “S.Score”, “G2M.Score” within the Seurat package. Clusters were identified using the FindClusters function by using Louvain resolution 1.2 for scRNA-seq.

Analysis, including data normalization, dimensionality reduction and clustering, of ST data were performed jointly using the Seurat and STUtility packages. Normalization was conducted using variance stabilizing transformation (SCTransform). Principal Component Analysis (PCA) was used for selection of significant components, a total of 50 principal components were used in downstream analysis and 30 principal components for ST analysis. To integrate ST sections Harmony (RunHarmony, version 1.0) function was used. Spots were clustered using the Shared Nearest Neighbor algorithm implemented in the Seurat package as FindNeighbors and FindClusters (Louvain resolution 0.7).

Uniform Manifold Approximation and Projection (UMAP) was used to create a 2D embedding of cell or spot transcription profiles for visualization purposes (RunUMAP). Identification of differentially expressed genes among clusters was done using the FindAllMarkers function from the Seurat package, where genes with log fold changes above 0.2 and p-values below 0.01 were considered significant. For the scRNA-seq data, a sub-selection of 50 random cells per cluster were used in order to compensate for differences in cell composition bias per cluster. For integration of scRNA-seq data and ST data we used stereoscope ^11^, which performs guided decomposition of the mixed expression data collected from each spatial capture location, using profiles learnt from scRNA-seq data as a reference. In the stereoscope analysis, a batch size of 2048 and 50000 epochs was used for both the parameter estimation step and the proportion inference process. Cell types with fewer than 25 cells were excluded from the analysis, while we randomly selected 500 cells from cell types with more than 500 members. For cell types with more than 25 members and less than 500 members, all cells were included. In the analysis, 2000 highly variable genes were used. These genes were extracted by applying the function scanpy.pp.highly_variable_genes() with n_top_genes=2000 from the scanpy (v.1.8.0.dev78+gc488909a) suite, after having normalized (scanpy.pp.normalize_total(…,target_sum=1e4)) and log-transformed (scanpy.pp.log1p(…)) the data. Cell type decomposition of ST spots was then saved as an assay for downstream analysis.

GO characteristics of gene clusters were determined using the clusterProfiler package (version 3.8.1) ^39^ for all DE genes with an average logFC value above zero, and an adjusted p-value below 0.01. The compareCluster function was used with a pvalueCutoff = 0.05. Analysis of genes belonging to Wnt, Shh or Notch pathways as well as Human Spinal Cord development were done using the KEGG database and Phenotype Orthologs (HPO), respectively.

For previously published scRNAseq data used in this study, data sources are listed below under the section Data availability. All these datasets were processed the same way as their publication stated.

#### Cell type annotation

After pre-processing and clustering analysis, each cluster (for both scRNA-seq and ST) was manually annotated based on previous knowledge and recent atlas resources. After annotating each cluster, clusters with the same major cell type names were merged and DEG analysis on these major cell types were performed in an unsupervised manner. These DEG results confirmed the accuracy of annotation. In addition, all available spinal cord scRNA-seq datasets (by June 2022) were integrated, and correlation analysis for annotations was performed, which showed high correlation between our dataset annotation and previous studies.

#### Inference of branching trajectories

The R package slingshot (version 1.8.0) ^15^ was used to analyze neurogenesis and gliogenesis _respectively._ For neurogenesis, the NPC cluster close to intermediate neuronal progenitors (INPs), all the INPs and all differentiated neurons were selected. For gliogenesis, we selected all glial cells and all NPCs that were connected to the trajectory. For each branch, clusters in the upstream and downstream were selected for pseudotime analysis. Lineage-associated genes were calculated by the R package TradeSeq (version 1.4.0) ^40^.

The R package URD (version 1.1.1) ^14^ was used to build differentiation trajectories during development. In the neurogenesis and gliogenesis analysis, a population of cells that were sampled from W5, clustered as NPCs_10, and with higher expression of *TOP2A* and *SOX2* was identified and used as root in the URD trajectory reconstruction. The tips of ASCs, OLs, EPCs, ExNs, ChNs and 3 IbNs lineages were identified based on the Louvain clusters (with a resolution 1.2), separately. After 350,000 simulated random walks were performed per tip, the divergence method “preference” was used to build the tree, with minimum.visits = 2, cells.per.pseudotime.bin = 25, bins.per.pseudotime.window = 8, p.thresh = 0.05 and min.cells.per.segment = 10.

In the inference of hNPC development trajectory, same population of NPCs was used as root, and the NPCs with later pseudotime estimated by scVelo and closer to neuronal and glial lineages on UMAP were identified as tips, respectively. The divergence method “preference” was also used for tree building, with cells.per.pseudotime.bin = 25, bins.per.pseudotime.window = 8, p.thresh = 0.001 and other parameters default.

#### Estimation of RNA velocities

The transcriptional dynamics of splicing kinetics were modelled stochastically with scVelo (version 0.2.4) ^13^ and projected onto the UMAP embedding as streamlines. To show the connectivity between different clusters, the transition probabilities of cell-to-cell transitions were estimated and projected onto the same UMAP embedding.

#### Inference of transcription factor activity

The SCENIC software (version 0.11.2) ^17^ was used to infer TF activities in human and mouse neural cells separately. In the human dataset, 10% cells in each subtype were randomly sampled and combined to infer gene regulatory network with GRNBoost2 algorithm. Then all neural cells were used to predict candidate regulons (cisTarget) and to estimate the cellular enrichment of the predicted regulons (AUCell). The top 5 regulons with the highest specificity in each cell type were selected using the regulon_specificity_scores() function implemented in Python. For each regulon, its activity in all cells was fitted and binarized to determine the “on” or “off” state, and further used to compute the “percent activated” in the Dot plots (Extended Data Fig. 10 and Supplementary Figure5).

The glial cells and neuronal cells were separately subset and the subtype specificity was recalculated within the subsets. Same analysis pipeline was applied to the mouse dataset, except that all mouse neural cells were used in the network inference step (instead of 10% random sampling).

#### Calculation of dorsal-ventral axis gene expression

In order to assess how certain feature values (e.g., gene expression or cell type proportions) varies along the dorsal-ventral (DV) axis, we designed a method to cast the 2D data into a different and more informative 1D representation relating to the aforementioned axis. More specifically, we sought to model the feature value as a function of the position along the DV-axis, that is yi = f(xi), where yi is the feature value of observation i while xi is the position of said observation on the DV-axis. Below we describe in detail how we obtained the values yi and xi as well as the character of the non-parametric function f.

First, to determine each observation’s position along the DV-axis, we had to define the DV-axis in each sample. Thus, we manually annotated all observations (spots) as either belonging to the ventral or dorsal region. We denoted the (mutually exclusive) sets of spots in the dorsal and ventral regions as D respectively V, we also let |.| represent the cardinality operator. Then, we selected a subset of observations (D’ and V’) of size min(|D|,200) respectively min(|V|,100) from each set, and computed the “DV-difference vectors” δs according to:

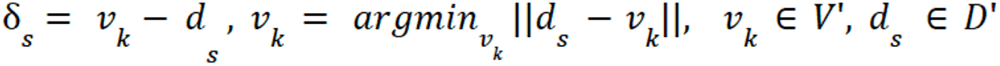

Whereafter we calculated the “average DV-difference vector”, representing the direction of the DV-axis, as follows:

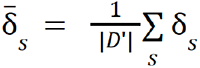

Finally, we let the axis vector a be defined as the normalized (to unit norm) average, across all observations within the sample, DV-difference vector. We then proceeded to project each observation’s spatial coordinates (in 2D space) onto the (1D) axis vector a, as to obtain its position along the DV-axis (ps); for this, standard orthogonal projection is used:

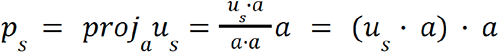

Where us is an observation’s original coordinates in the 2D plane, the final equality holds true since a has unit norm. For each sample, we then normalized the axis projections using min-max scaling (subtraction of minimal value and division with the difference between maximal and minimal values). For computational reasons, we assign each observation s (based on their axis projection value) to one bin (bi) of nbins different bins, according to:

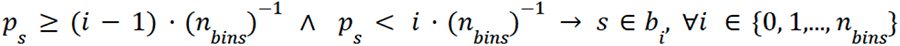

Next, for each bin bi we compute the average axis value (xi) and average feature value (yi) as follows:

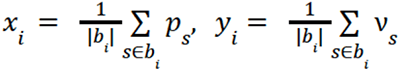

Where νs is the feature value associated with observation s. In the last step, we aim to relate the feature values to the axis positions via a function f. The character of f is determined by loess regression (locally estimated scatterplot smoothing), implemented with geom_smooth(…, method = loess) from the R package ggplot2 and visualized as a 1D plot - generating the plots similar to those shown (for example) in Figure 3h.

We implemented this method in R, and all code is available at GitHub (https://github.com/almaan/axis-projection) as a package that can be installed and used in a standard R environment.

#### Image processing and decoding for HybISS data

After imaging, Leica LAS X software was used to maximum intensity project each field of view (FOV) to obtain a flattened two-dimensional image. Imaging data was then analyzed with in-house custom software that handles image processing and gene calling based on the python package Starfish. Each two-dimensional FOV was exported, and preprocessed by alignment between cycles, and stitched together using the MIST algorithm ^41^. Stitching was followed by retiling to create smaller non-overlapping 2000×2000 pixel images that were then used for decoding. The decoding pipeline can be found on the Moldia GitHub page (https://github.com/Moldia/iss_starfish/). In short, the images were initially registered (using the LearnTransform module in Starfish) and filtered (using the Filter module from Starfish) using a white top hat filter with a masking radius of 15. The filtered images are subsequently normalized (using the Filter module from Starfish). Following the normalization, spots were detected using the FindSpots module from Starfish and subsequently decoded using MetricDistance decoding.

#### Probabilistic cell typing for HybISS data

Probabilistic cell maps were created using probabilistic cell typing by in situ sequencing (pciSeq). The pciSeq pipeline can be found at https://github.com/acycliq/pciSeq and is described in Qian et al. ^12^ In short, pciSeq works by assigning genes to cells and then cells to cell types, and this assignment is done using a probabilistic framework based on a single-cell RNA sequencing data ^12^. Due to the density of nuclei in the tissue, nuclear segmentation could not be done, instead a compartment-based approach was employed in which each compartment was defined as 40×40 pixel grid (roughly 13×13 µm).

### Quantification and Statistical Analysis

Significance of scRNA-seq and ST analysis for differential gene expression were carried out using Wilcox. Genes with P < 0.001 were selected as significantly different expressed genes. Significantly different expressed gene lists were ordered and filtered by smallest P value the biggest changes of log Fc.

## Data and Code Availability

Codes for analysis of this paper can be found from the link: https://github.com/czarnewski/human_developing_spinal_cord

Data will be made publicly available on Gene Expression Omnibus (GEO) upon publication. The publicly available data utilized in this study are available at:

Sathyamurthy: https://www.ncbi.nlm.nih.gov/geo/query/acc.cgi?acc=GSE103892

Zeisel: https://www.ncbi.nlm.nih.gov/sra/SRP135960

Rosenberg: https://www.ncbi.nlm.nih.gov/geo/query/acc.cgi?acc=GSE110823

Blum: https://www.ncbi.nlm.nih.gov/geo/query/acc.cgi?acc=GSE161621

Alkaslasi: https://www.ncbi.nlm.nih.gov/geo/query/acc.cgi?acc=GSE167597

Delile: https://www.ebi.ac.uk/arrayexpress/experiments/E-MTAB-7320/files

Rayon: https://www.ncbi.nlm.nih.gov/geo/query/acc.cgi?acc=GSE171892

Milich: https://www.ncbi.nlm.nih.gov/geo/query/acc.cgi?acc=GSE162610

Zhang: https://www.ncbi.nlm.nih.gov/geo/query/acc.cgi?acc=GSE136719

Gojo (ependymomas): https://www.ncbi.nlm.nih.gov/geo/query/acc.cgi?acc=GSE141460

## Acknowledgements

The study was supported by the Erling Persson Family Foundation, Knut and Alice Wallenberg Foundation and the research funds of Karolinska Institutet and Science for Life Laboratory. P.C. is partly financially supported by the Knut and Alice Wallenberg Foundation as part of the National Bioinformatics Infrastructure Sweden at SciLifeLab. The National Genomics Infrastructure (NGI), Sweden is acknowledged for providing infrastructure support. Chan Zuckerberg Initiative, an advised fund of Silicon Valley Community Foundation [2018- 191929] and the Swedish Research Council [2019-01238] to M.N.; Swedish Brain Foundation (Hjärnfonden) [PS2018-0012] to D.G; Chinese Scholarship Council to Y.L. The authors acknowledge the support by the Karolinska Institutet Developmental Tissue Bank for providing human fetal tissues.

## Author information

X.L. designed the study, planned, and performed the experiments (human tissue dissection, scRNA-seq, ST and HybISS), analyzed the scRNA-seq data, interpreted bioinformatic results (scRNA-seq, ST, HybISS and IHC) and biological results, wrote the manuscript, and designed and prepared figures. Za.A. performed ST experiments, analyzed the ST data, interpreted results, and designed figures. P.C. guided bioinformatic data analysis, interpreted results and designed figures. A.A. conducted the stereoscope analysis and developed the method to examine feature values along the DV-axis. C.M.L. analyzed the HybISS data and prepared figures. Y.L. participated in trajectory analysis. D.G. performed HybISS experiments and guided HybISS data analysis. E.B. performed scRNA-seq experiments. L.L. guided data analysis for ST. L.H. supported scRNA-seq experiments. Zh.A. performed IHC. H.K.K. managed the collection of prenatal human tissue. E.Å. dissected and staged the tissue material. M.N. guided HybISS experiments. S.L. guided scRNA-seq experiments. I.A. participated in experimental and data analysis discussion. J.L. guided ST experiments. E.S. conceived and designed the study, dissected, and staged the tissue materials, provided biological guidance, supported biological result interpretation, and wrote the manuscript. All authors helped with manuscript preparation.

## Ethics declarations

Z.A., L.L., J.L. are consultants, and M.N. advisor for 10x Genomics Inc.

## Figure Legends

**Extended Data Fig. 1.**
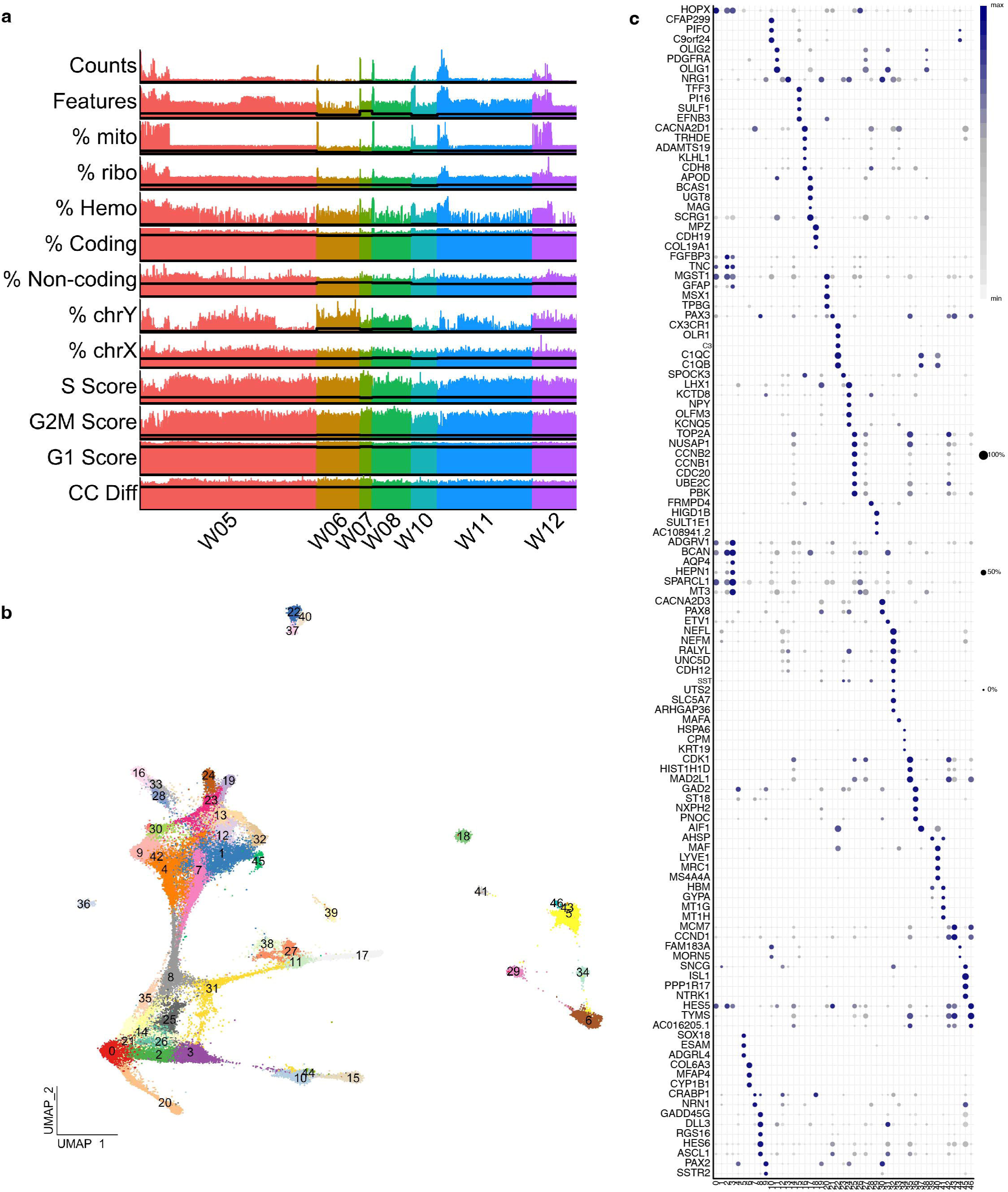
scRNA-seq reveals cell heterogeneity of the developing human spinal cord. a) Quality control and filtering strategies for scRNA-seq dataseq. Thick lines indicate filter thresholds. b) UMAPs identifying 47 clusters (cluster 0-46). c) Dot plot illustrating top marker genes of each cluster. In relation to Fig 1.

**Extended Data Fig. 2.**
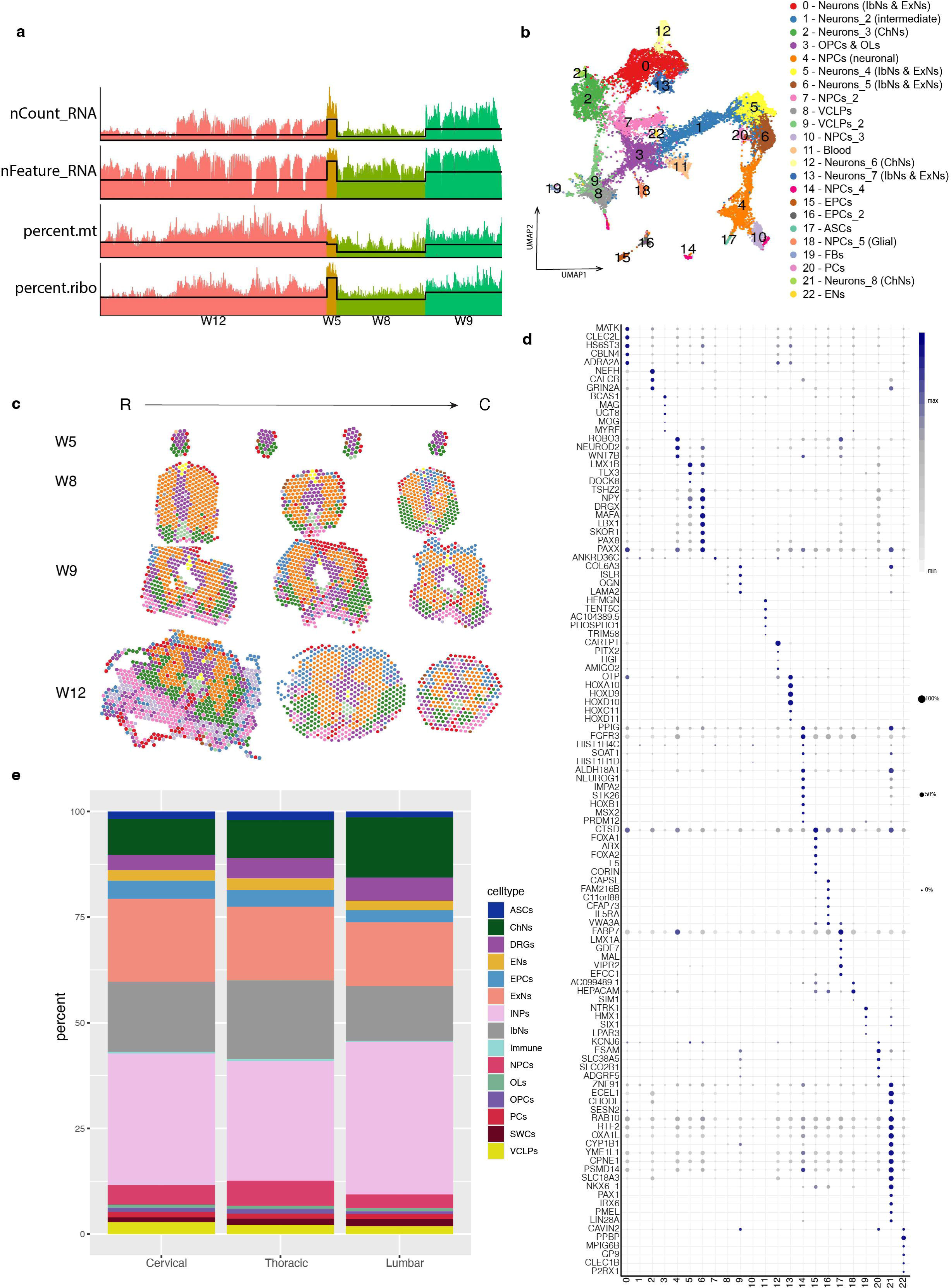
Temporal and spatial gene expression in the developing human spinal cord by ST. a) Quality control results of ST. Lines indicate filtering thresholds. b) UMAP illustrating 23 clusters from 76 ST sections at W5, W8, W9 and W12. C) Representative sections of ST spatial maps of all clusters along rostral-caudal axis. R= rostral, C = caudal. D) Dot plot illustrating the top marker genes for all clusters in the ST analysis. e) Bar graph illustrating cell type proportions across sections along rostral-caudal axis. In relation to Fig 1.

**Extended Data Fig. 3.**
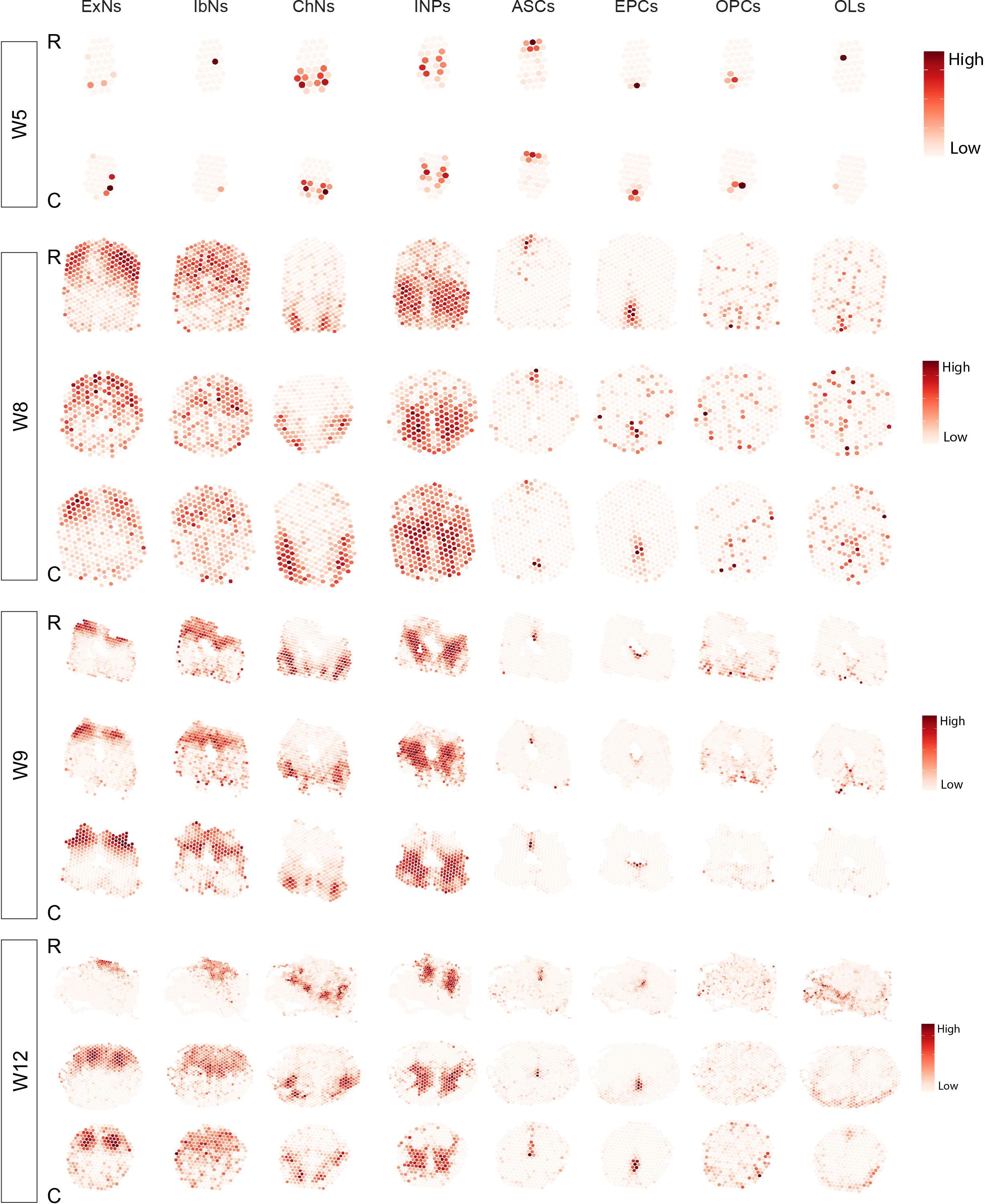
Stereoscope revealing integrated results of scRNA-seq and ST. Representative stereoscope images revealing cell type positions and their probability along the rostral-caudal axis at W5, 8, 9 and 12. R = rostral, C = caudal. Intermediate neuronal progenitors (INPs), excitatory neurons (ExNs), inhibitory neurons (IbNs), cholinergic neurons (ChNs), astrocytes (ASCs), ependymal cells (EPCs), oligodendrocyte precursor cells (OPCs), oligodendrocytes (OLs). In relation to Fig 1.

**Extended Data Fig. 4.**
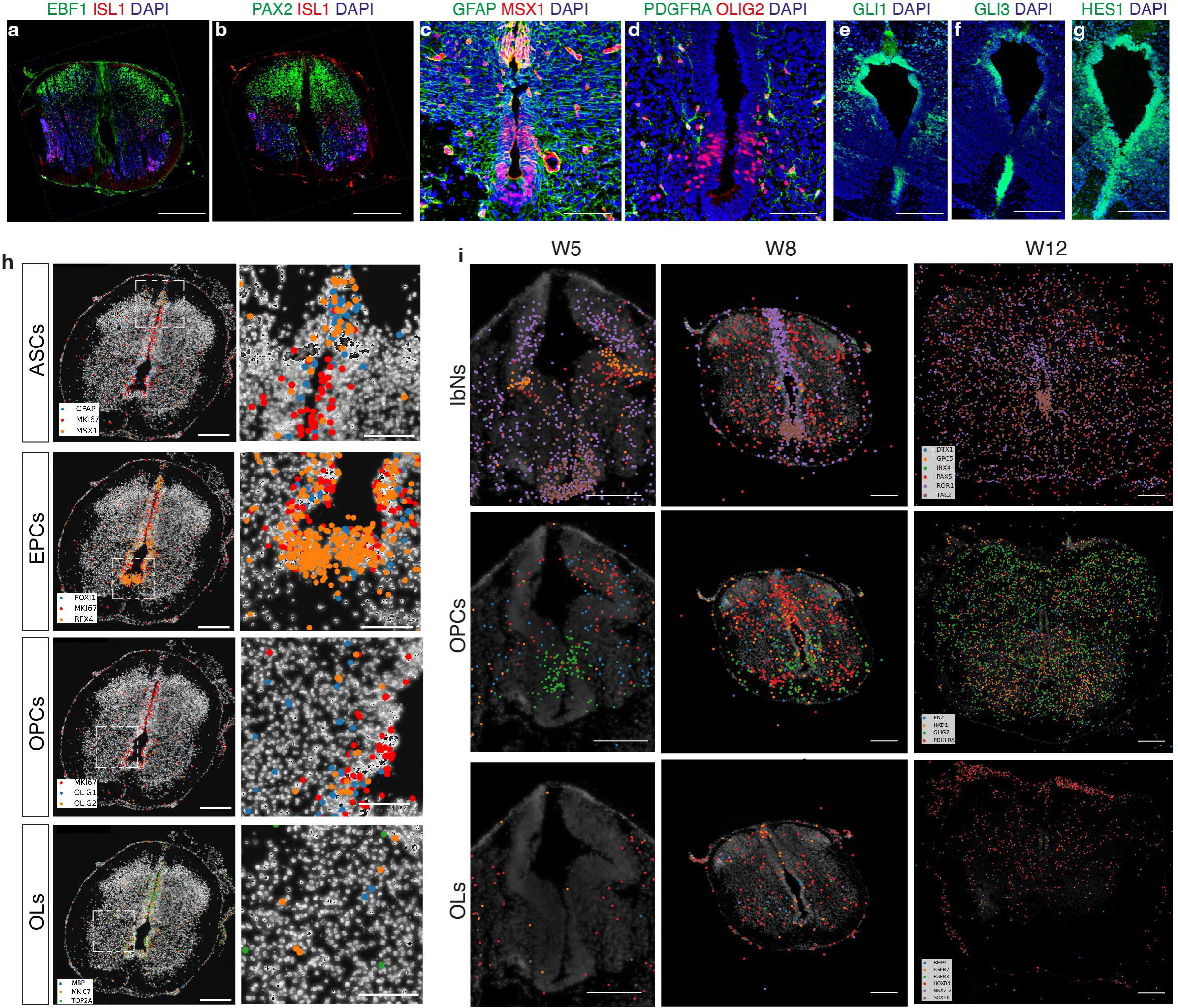
Validation of cell populations, cell fate commitment and neural patterning at early developmental stage. a-g) Representative IHC images illustrating localization of ExNs, IbNs and ChNs (a-b), ASCs (c) and OPCs (d) in W8 human spinal cord as well as SHH and Notch related proteins at W5 human spinal cord sections (e-g). h) Representative HybISS images illustrating early glial cells at W8. Rectangles indicating enlarged areas. i) Representative HybISS images illustrating localization of subpopulations of IbNs, OPCs and OLs during human spinal cord development. Scale bars: 100 μm. In relation to Fig 2 and Fig 4.

**Extended Data Fig. 5.**
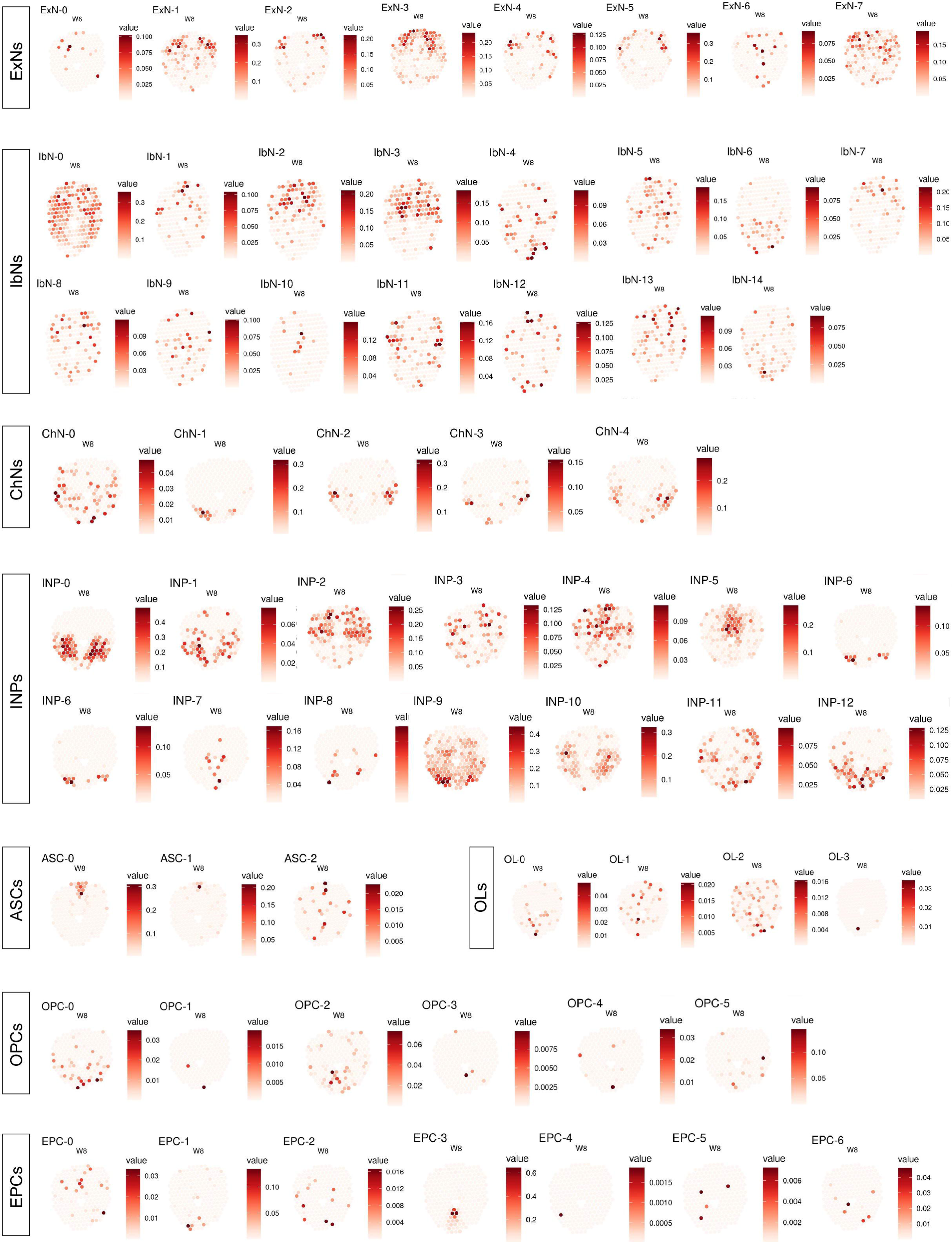
Locolization of heterogenous cell types or cell states of neural cells in the human developing spinal cord. Representative images from stereoscope analysis illustrating the probability of spatial distribution of each cell subpopulation or cell state of the major neural cells at W8. In relation to Fig 2.

**Extended Data Fig. 6.**
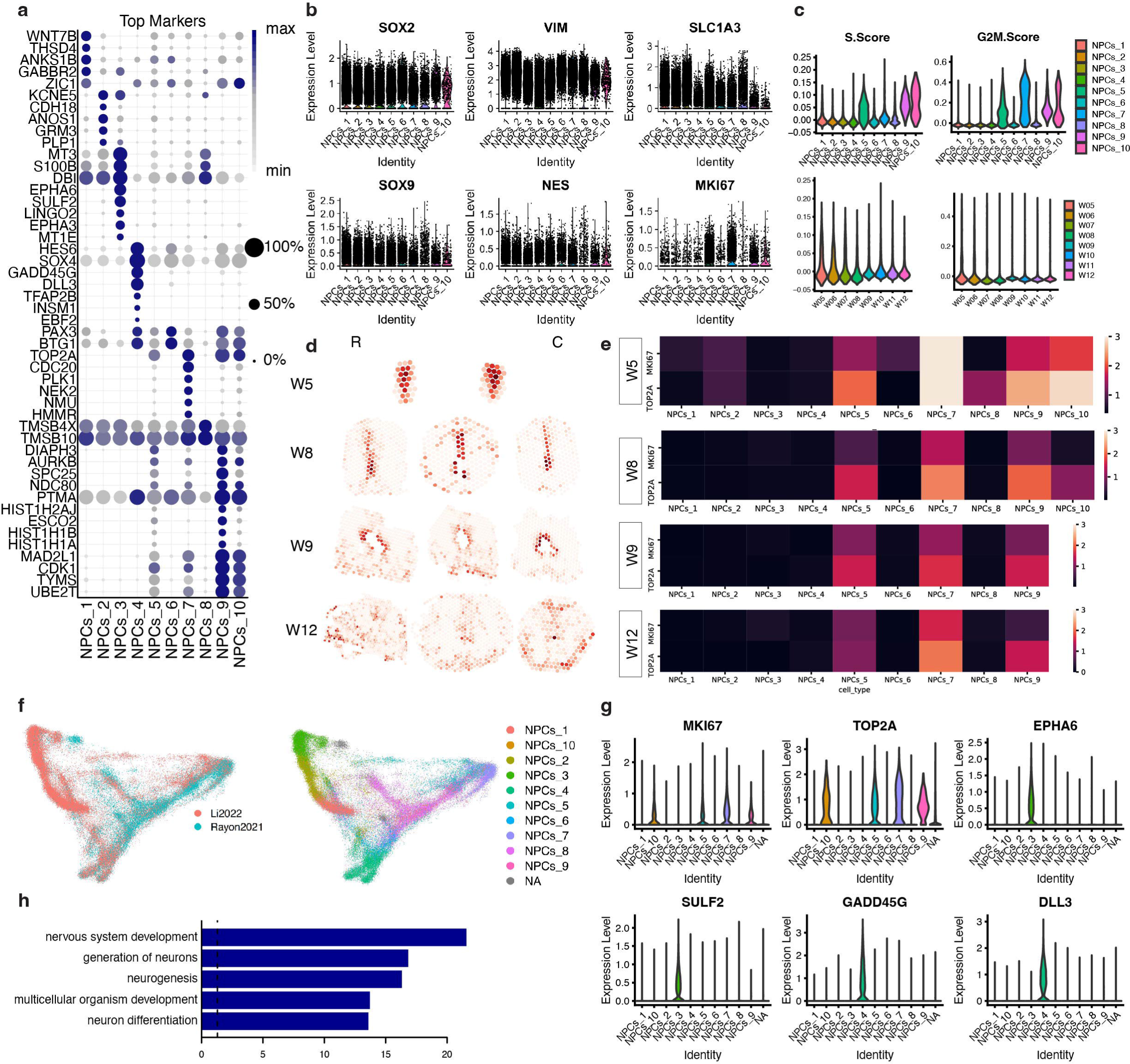
Heterogenous NPCs with different activeness in the developing human spinal cord. a) Dot plot illustrating the top markers for the NPC clusters. b-c) Violin plots illustrating stem cell markers (b) and cell cycle scores (c) of human NPCs across clusters and ages. d) Stereoscope illustrating the probability of spatial distribution of different NPCs in the developing human spinal cord sections. e) HybISS illustrating the locations of the proliferative NPC clusters from W5-12. Scale bar 100μm. f) UMAP illustrating integrated datasets and subtypes of NPCs. g) Violin plots illustrating consistent results for gene expression of proliferation markers and subtype specific markers in the integrated dataset. h) Top GO terms of early non-proliferative NPCs compared to proliferative NPCs. In relation to Fig 2-3.

**Extended Data Fig. 7.**
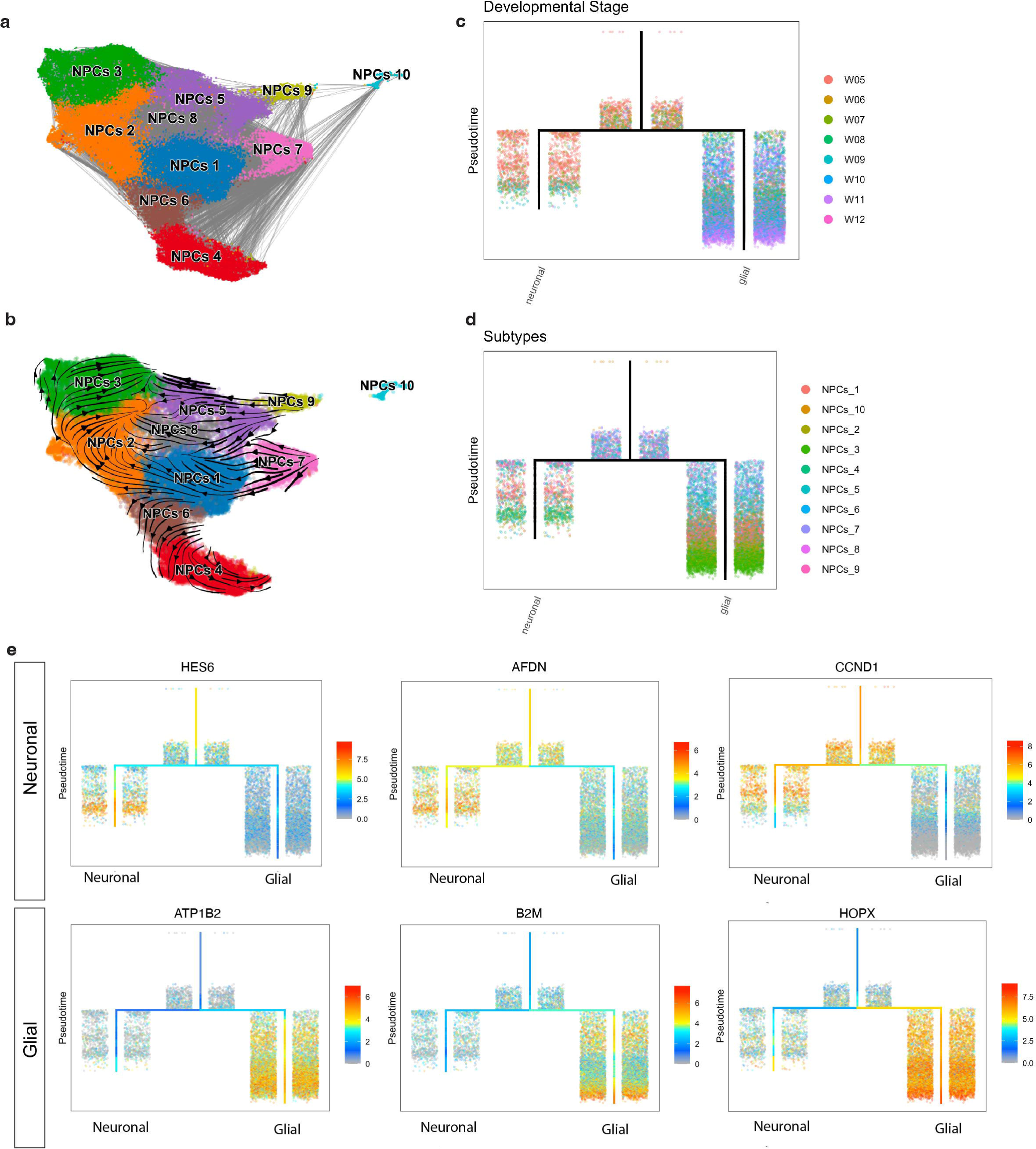
Cell fate commitment of hNPCs during human spinal cord development. a) UMAP showing strong connectivity of different NPC clusters during development. b) scVelo analysis revealing the predicted differentiation trajectory from proliferative NPCs to neuronal and glial fate committed NPCs. c-d) Hierarchical tree from URD analysis displaying NPC trajectory during development. e) Hierarchical trees illustrating examples of top genes associated with neuronal and glial lineage during cell fate commitment of hNPCs. In relation to Fig 3.

**Extended Data Fig. 8.**
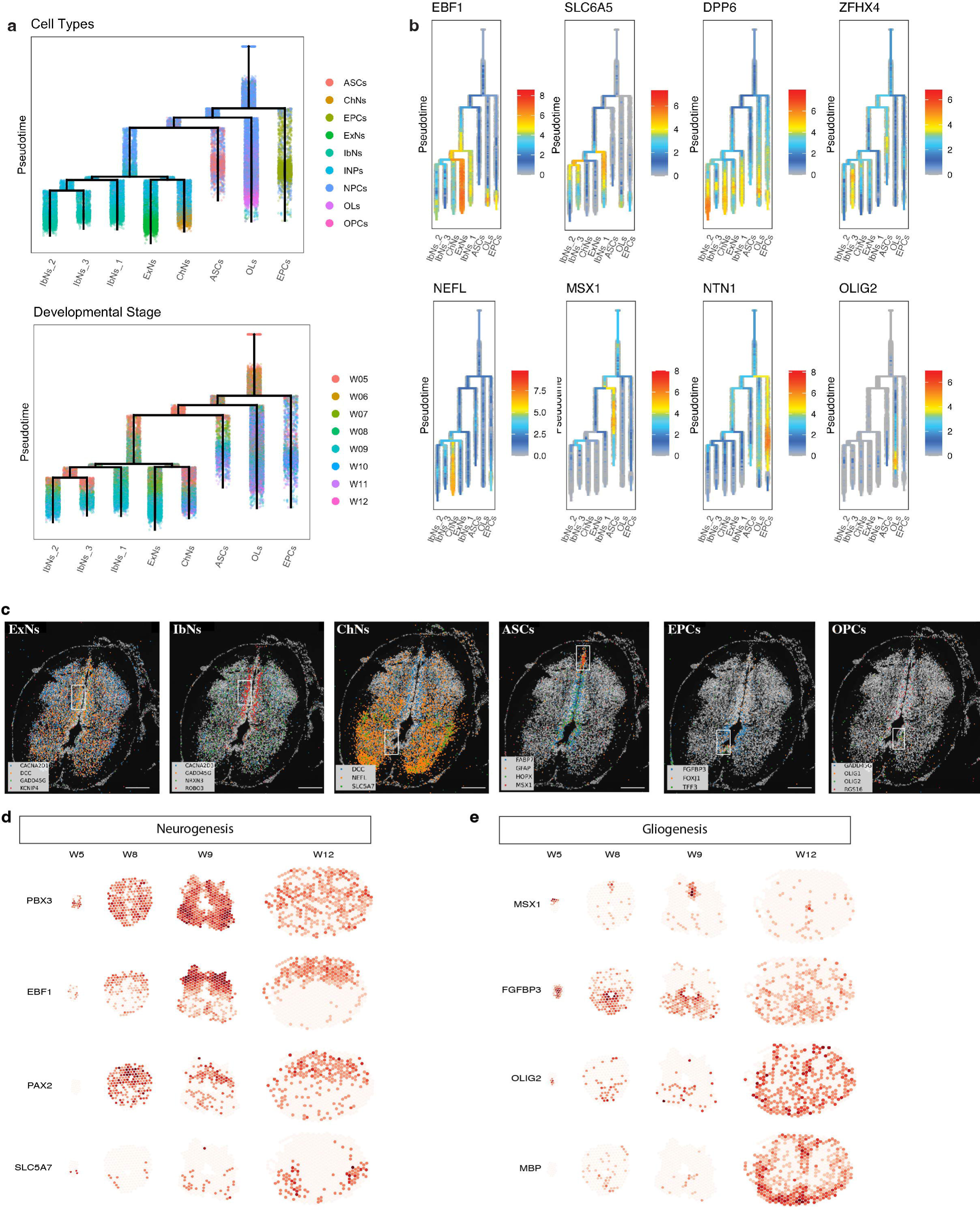
Spatiotemporal regulation of human neurogenesis and gliogenesis. a-b) Hierarchical trees from URD analysis displaying neurogenesis and gliogenesis (a) as well as top lineage-associated genes (b) during human spinal cord development. c) Representative HybISS images illustrating the co-locolization of NPC markers, committed cell fate markers and lineage-associated genes. Scale bar 100μm. e) ST plots illustrating the spatial expression of top lineage genes revealed by scRNA-seq. In relation to Fig 3.

**Extended Data Fig. 9.**
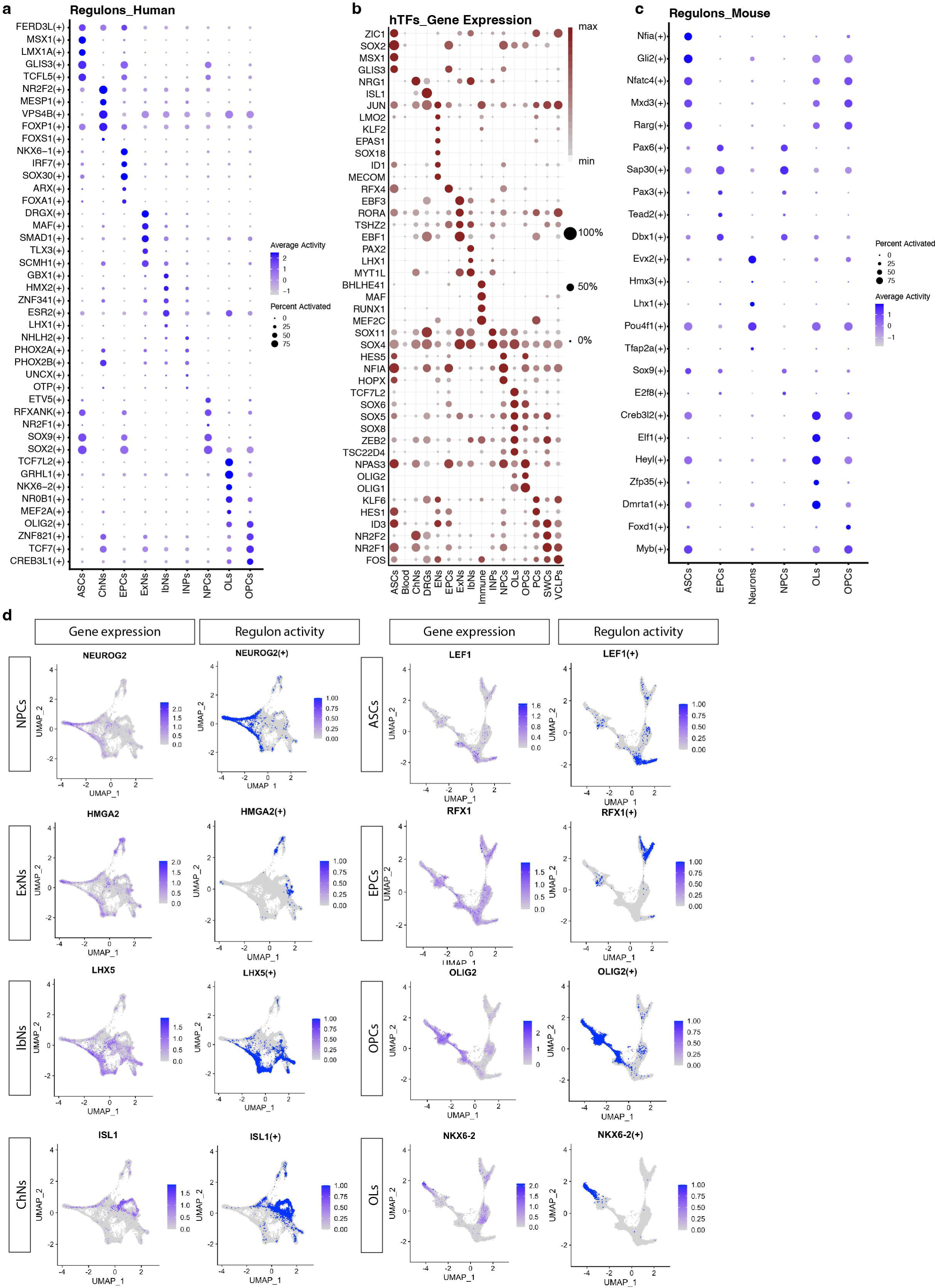
Lineage associated regulons have species difference during neurodevelopment. a-b) Dot plots illustrating top regulons (a) and their gene expression (b) in human major cell types during development. c) Dot plot illustrating top regulons during mouse spinal cord development. d) Dot plots illustrating top regulons and their gene expression during human neurogenesis and gliogenesis. In relation to Fig 5.

**Extended Data Fig. 10.**
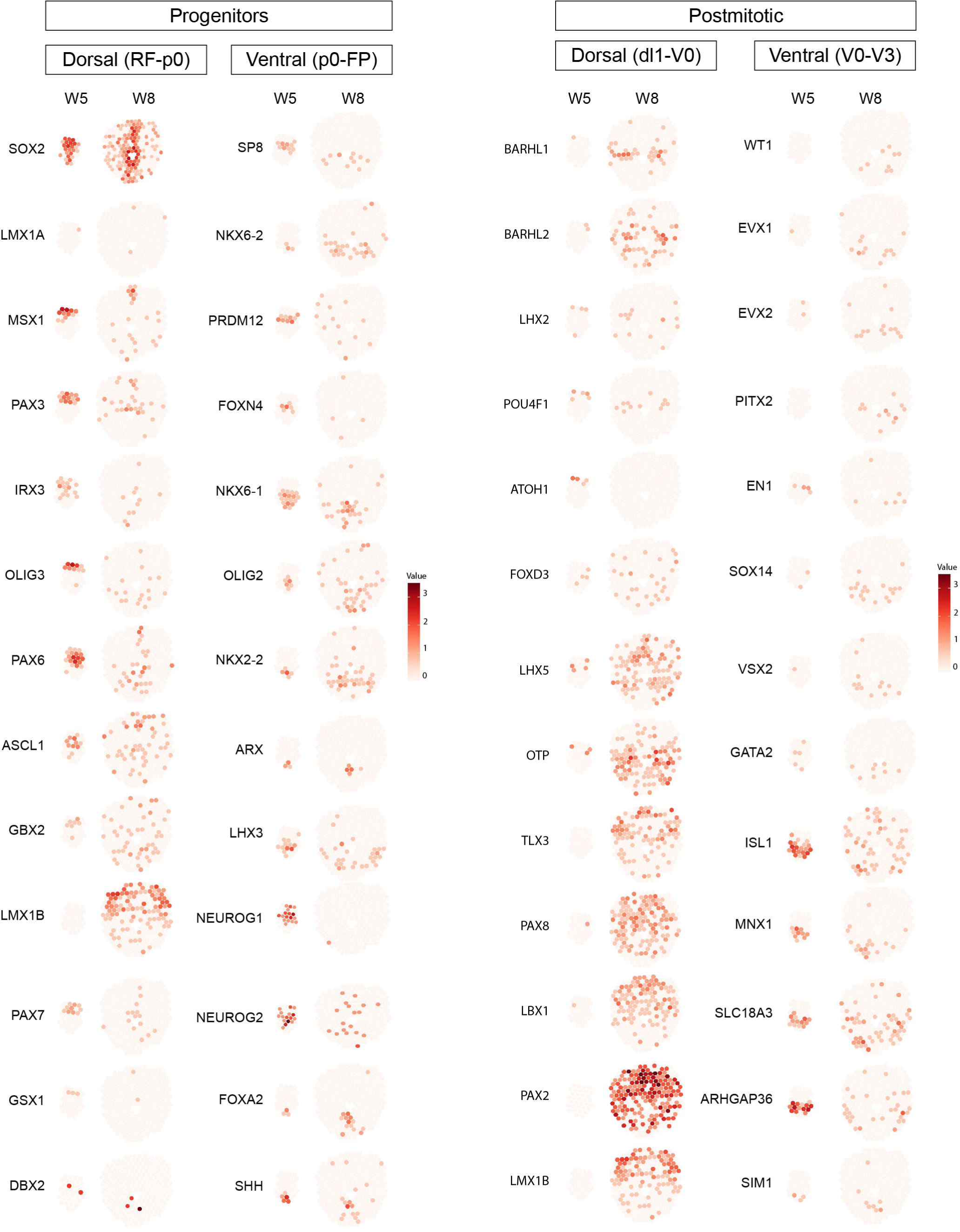
Neural patterning gene expression in the early developmental human spinal cord. Neural patterning genes related to progenitor and postmitotic neurons are plotted in W5 and W8 representative human spinal cord sections along the dorsal-ventral axis. Most of the neural patterning genes enriched in the progenitors appear at W5 but most of them disappear at W8 (Left panel). In contrast, neural patterning genes expressed in neurons are mostly absent at W5 but exhibit a dorsal-ventral patterns at W8. In relation to Fig 4.

**Supplementary Figure 1. Selection of probes for HybISS based on major cell type markers.** Dot plot illustrating the correlation between major cell types from scRNA-seq and the chosen probes for validation with HybISS.

**Supplementary Figure 2. Selection of probes for HybISS based on subpopulations of each major cell type markers.** Dot plot illustrating the correlation between subtypes or cell states within each major cell type from scRNA-seq and the chosen probes for validation with HybISS.

**Supplementary Figure 3. Spatiotemporal gene expression regulates neurogenesis and gliogenesis**. a) Dot plot illustrating different markers among three lineages of IbNs. b) GO terms of three terminal clusters of IbNs. c) Minimal spanning tree (MST) displaying the strongest connections between clusters related to neurogenesis. d) Heatmaps illustrating lineage differential gene expression of each branch. e) MST displaying the strongest connections between clusters related to gliogenesis. f) Heatmaps illustrating lineage differential gene expression of each branch.

**Supplementary Figure 4. The regulatory networks of human spinal cord development.** a) Dot plot illustrating the expression of signaling pathway genes in all major cell types throughout W5-12 developmental stages. b) Interactome analysis indicate the interaction between every two cell types and the ligands and receptors contributing to the interactions. L = ligand, R = receptor. Red lines indicate increased expression of related ligands or receptors, while blue lines indicate decreased expression of related ligands and receptors during development. Thicker lines between ligands and receptors indicate higher probability of connections and cell-cell interactions.

**Supplementary Figure 5. Top regulons during human and mouse spinal cord development.** a) Dot plot illustrating the most significant regulons in the developing human spinal cord during W5-12 across major cell types and age. b) Dot plot illustrating the most significant regulons in the developing mouse spinal cord from E9.5-P11 across major cell types and age.

**Supplementary Figure 6. Fetal human spinal cord and relation to ependymomas.** a) UMAP displaying all major cell types revealed by the integrated scRNA-seq dataset of human developing spinal cord and human ependymomas. b) Heatmaps revealing the most significantly differential expressed genes in two lineages of EPCs-like cells and ASCs and OPCs/OLs-like cells during trajectory analysis. c) Dot plot illustrating the expression of putative cancer stem cell marker genes across all subtypes of human NPCs and human CSCs and is selectively enriched in cluster 3, 6, and 7.

**Supplementary Figure 7. Correlation with annotations from previous spinal cord datasets.** Heatmaps illustrating good correlation between the annotation of scRNA-seq data from this study with previous scRNA-seq data by two different correlation calculations.

